# Developmental deviations of structural connectivity in youths with ADHD predict symptom and treatment outcomes

**DOI:** 10.1101/2025.11.20.689391

**Authors:** Xiaoyu Xu, Zhao Fu, Haoshu Xu, Kangfuxi Zhang, Hang Yang, Yang Li, Yilu Zhao, Xiangyu Zheng, Arielle Keller, Ting Xu, Runsen Chen, Weidong Cai, Qingjiu Cao, Li Yang, Zaixu Cui

**Author notes:** Correspondence (Z.C.), (L.Y.), (Q.C.). These authors contributed equally to this work (Xiaoyu Xu, Zhao Fu).

## Abstract

Attention-deficit/hyperactivity disorder (ADHD) lacks validated biomarkers to track symptom development or guide treatment selection in youth. We quantified individual deviations from normative age-related trajectories of white-matter structural connectivity (SC) across two independent cohorts: 6,687 typically developing and 1,114 ADHD scans from the ABCD study, and 292 typically developing and 482 ADHD participants from a Chinese cohort. Youth with ADHD showed pronounced SC deviations, especially in higher-order association pathways. A subset of connections showed ADHD-specific developmental reductions in deviations beyond typical developmental patterns, and these changes mediated the age-related decline in ADHD symptoms. Longitudinal within-individual analyses demonstrated that decreases in deviation over two years tracked symptom improvement. Baseline deviations also predicted 12-week treatment response to atomoxetine, but not methylphenidate, and follow-up imaging revealed treatment-related reductions in deviation. Together, these findings identify SC deviation as a robust developmental biomarker with prognostic and theragnostic relevance for ADHD, supporting precision care through risk stratification, individualized pharmacotherapy selection, and objective monitoring of treatment effects.

## Introduction

Attention-deficit/hyperactivity disorder (ADHD) is a prototypical neurodevelopmental disorder that emerges in childhood and undergoes marked changes during youth^1^. It affects approximately 5% of school-aged children worldwide and is characterized by symptoms of inattention and hyperactivity-impulsivity that evolve across development^1^. While some individuals experience remission during youth, many remain symptomatic into adulthood^2,3^. Treatment response is similarly heterogeneous, with patients of comparable symptom severity showing divergent outcomes to the same medication^4,5^. This variability in prognosis and treatment effects likely reflects underlying neurodevelopmental processes that are not captured by clinical manifestation alone^6–8^. Identifying brain-based markers that characterize these developmental processes, particularly during youth as a period of heightened neuroplasticity, could therefore improve prediction of both symptom trajectories and treatment response.

Converging evidence from structural and functional MRI has revealed abnormalities across distributed regions spanning the frontal, parietal, and temporal cortices^9,10^, which collectively involved in attention, cognitive control, and motivation. Structural connectivity (SC), defined by white-matter pathways forming the anatomical backbone for inter-regional communication^11^, provides a unifying substrate linking these distributed abnormalities in ADHD. SC undergoes protracted maturation through myelination and axonal remodeling, with higher-order association pathways developing most slowly and remaining most plastic^12^. Such prolonged plasticity may confer both vulnerability to atypical wiring and opportunities for adaptive change. Prior neuroanatomical work has shown delayed cortical maturation in ADHD, particularly within prefrontal regions supporting executive control^13,14^. However, whether white-matter SC follows a similar deviation from normative developmental trajectories remains unclear.

Diffusion MRI (dMRI) studies have reported white-matter abnormalities in ADHD, yet findings remain heterogeneous^15^. A recent meta-analysis of dMRI studies revealed no robust microstructural alterations in white matter among children and adolescents with ADHD^15^. This inconsistency may reflect the limitations of traditional case-control designs, which compare static group means and overlook that normative white-matter organization changes across childhood and adolescence, resulting in age-dependent shifts in developmental benchmarks^16^. A recent normative-modeling study began to move beyond such contrasts by quantifying individual deviations from typical white-matter development in ADHD^17^. However, that work focused on predefined tracts, such as the superior longitudinal fasciculus, which did not specify the cortical regions they connected to^17^. Whole-brain connectome approaches address this limitation by quantifying SC between functional systems, offering a more comprehensive and functionally interpretable framework for understanding the connectivity basis of ADHD. Moreover, prior work^17^ was based on a modest sample size and the lack of longitudinal or treatment data, limiting its ability to capture developmental trajectories and to assess the effects of clinical interventions.

To address these gaps, we combined large-scale longitudinal and cross-cultural diffusion MRI data to map normative developmental trajectories of cortico-cortical structural connectivity and identify ADHD-related deviations during youth. Using the Adolescent Brain Cognitive Development (ABCD) study^18^ as a discovery cohort and an independent Chinese cohort for replication and treatment analyses, we tested three hypotheses: first, that ADHD is characterized by deviations concentrated in higher-order association pathways; second, that these deviations track developmental changes in ADHD symptoms; and third, that they predict differential treatment response. By integrating developmental, cross-cultural, and pharmacological data, this study identifies SC deviations as developmental markers of ADHD with translational potential for guiding individualized treatment strategies.

## Results

We leveraged a longitudinal discovery dataset from the ABCD study^18^ (baseline, 2-year, and 4-year follow-ups) and an independent cross-sectional dataset from a Chinese cohort with treatment follow-up. The ABCD dataset included 6,687 scans from typically developing (TD) children and 1,114 from children with ADHD (ages 8.9–15.5) (**Fig. 1A**), while the Chinese cohort comprised 292 TD and 468 ADHD participants (ages 6.5–15.5 years) (**Fig. 1B**). Inclusion and exclusion criteria and full demographics are provided in **Fig. S1–S2** and **Tables S1–S2**.

**Fig. 1.**
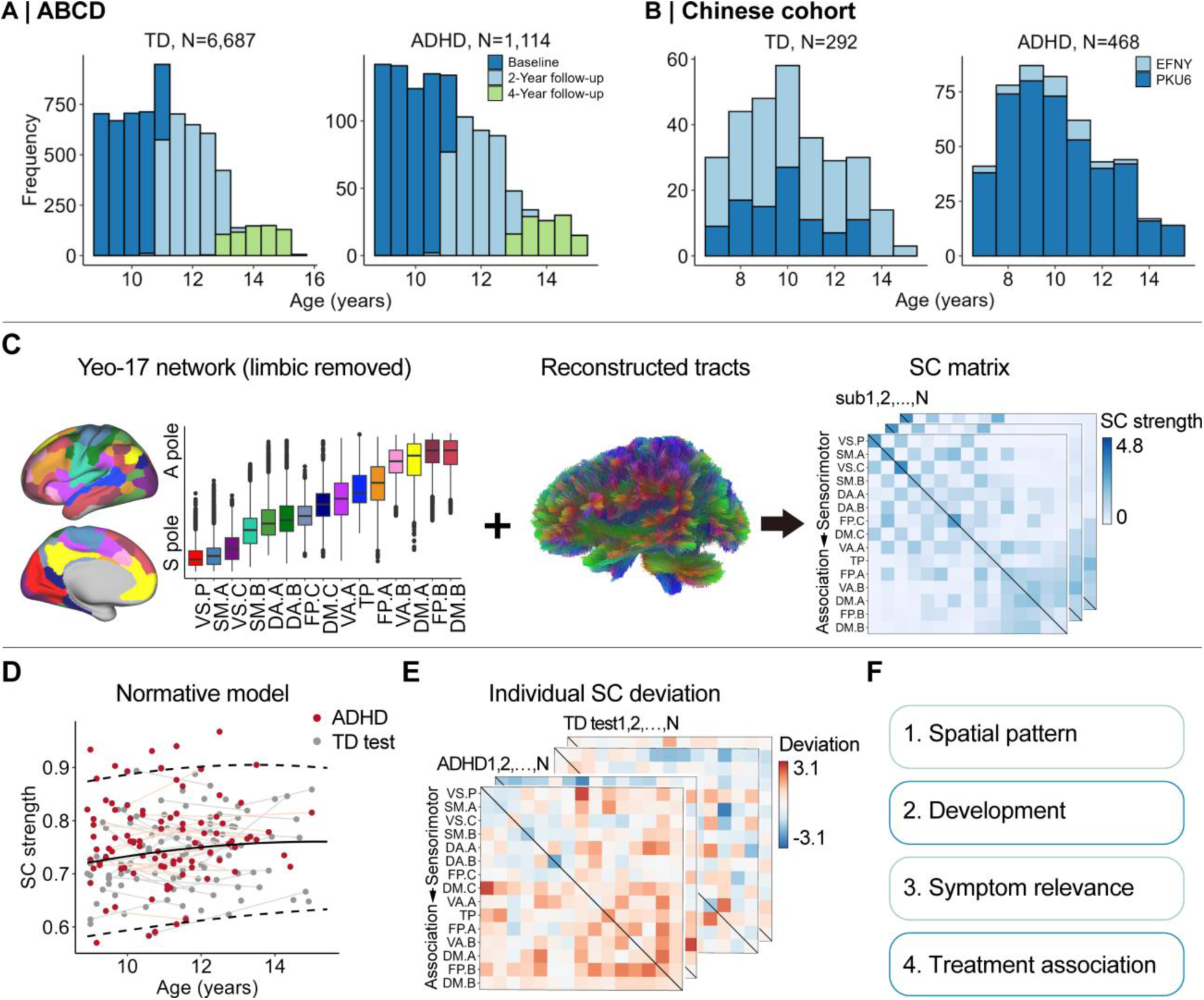
Overview of the study framework. **A**, Age distribution of participants in the ABCD discovery cohort. **B**, Age distribution of participants in the Chinese cohort. **C**, Cortical networks were defined using the Yeo-17 parcellation, with limbic networks excluded^23^, and ordered by their position along the S-A cortical axis^20^. For each scan, whole-brain white matter tracts were reconstructed from diffusion MRI, and a 15 × 15 SC matrix was derived. Each matrix contained 120 undirected connections, with connection weights defined as streamline counts between network pairs normalized by their cortical volumes^24^. **D**, Developmental normative models of SC strength were estimated for each connection using a training subset of TD youths. **E**, For each participant and visit, individual SC deviation scores were computed relative to the age-expected normative trajectories for both the TD test and ADHD groups. **F**, SC deviation maps were analyzed to characterize their spatial organization, developmental changes, and associations with clinical symptoms and medication effects in youths with ADHD. VS.C: Visual central; VS.P: Visual peripheral; SM.A: Somatomotor A; SM.B: Somatomotor B; DA.A: Dorsal attention A; DA.B: Dorsal attention B; VA.A: Salience/Ventral attention A; VA.B: Salience/Ventral attention B; FP.C: Frontoparietal control C; FP.A: Frontoparietal control A; FP.B: Frontoparietal control B; TP: Temporal parietal; DM.C: Default mode C; DM.A: Default mode A; DM.B: Default mode B; SC: structural connectivity; S-A: sensorimotor-association; ADHD: attention-deficit/hyperactivity disorder; TD: typically developing; ABCD: Adolescent Brain Cognitive Development; EFNY: Executive Function and Neurodevelopment in Youth; PKU6: Peking University Sixth Hospital.

For each scan, we reconstructed a cortico-cortical structural connectivity (SC) matrix from diffusion MRI (dMRI) and T1-weighted imaging (T1WI) (**Fig. 1C**). Cortical systems were defined by the Yeo-17 network parcellation, excluding the limbic networks due to low signal-to-noise ratio^19^. The remaining 15 networks were arranged along the sensorimotor-association (S-A) cortical axis^20^, which is a canonical hierarchical gradient progressing from primary sensorimotor to higher-order association regions, to facilitate visualization and interpretation of connectome-wide developmental patterns.

Normative developmental models for each of the 120 inter- and intra-network connections were trained using generalized additive models for location, scale, and shape (GAMLSS)^21,22^ (**Fig. 1D**). To ensure robust estimation and independent validation, the TD sample from the ABCD study was randomly divided into a training set (two-thirds) and a held-out test set (one-third), stratified by assessment wave. The TD training set was used to estimate age-dependent normative trajectories of SC strength for each connection. For each participant at each visit, connection-wise deviation scores were computed in both the TD test set and the ADHD group, quantifying how individual SC strength deviated from age-expected norms (**Fig. 1E**). We further examined the connectome-wide spatial organization, developmental dynamics, symptom relevance, and treatment associations of these connectivity deviations in children with ADHD (**Fig. 1F**).

### Delayed maturation of structural connectivity in ADHD

Using the discovery dataset from ABCD study, we modelled the developmental trajectories for each of the 120 connections separately in the TD and ADHD groups using GAMLSS, with age, sex, and head motion as covariates and site as a random effect (See Methods). Across late childhood to mid-adolescence, most connections strengthened with age in both TD (**Fig. 2A**) and ADHD (**Fig. 2B**) groups. To quantify developmental change, we calculated the mean age-related slope (first derivative) for each connection (**Fig. 2C**). In both groups, connections involving higher-order association networks, particularly those linking frontoparietal and default mode networks, showed smaller and later increases than sensorimotor connections. However, these age-related slopes were weaker in ADHD, indicating attenuated strengthening of association-related pathways.

**Fig. 2.**
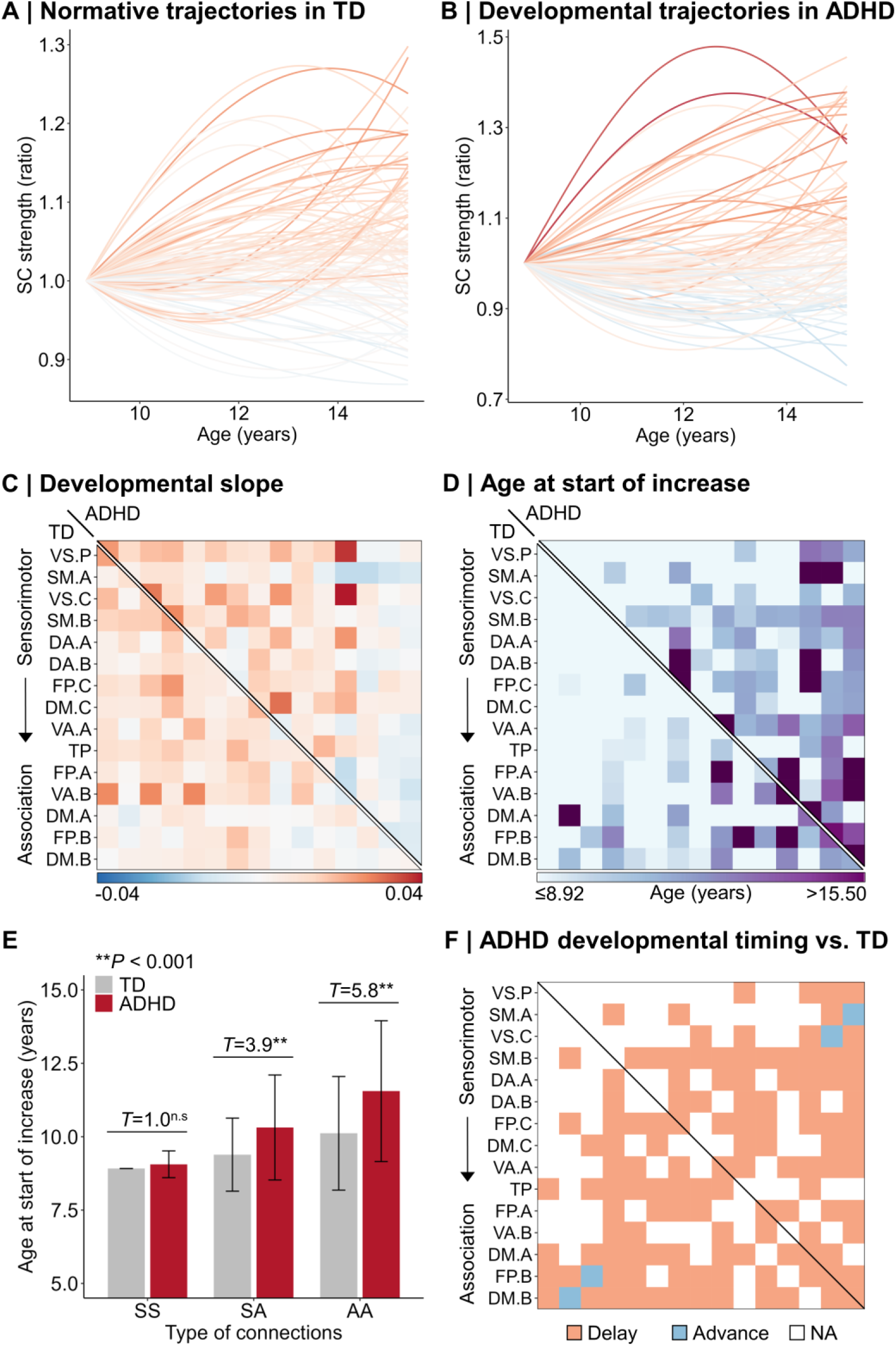
Delayed maturation of structural connectivity in ADHD. **A**, Normative developmental trajectories of SC strength across 120 connections in TD youths. **B**, Developmental trajectories of SC strength in youth with ADHD. Each curve represents the median trajectory for one connection, color-coded by its mean first derivative, as shown in (**C**). **C**, Average first derivatives representing developmental slopes for all connections. **D**, Ages at which SC strength began to increase, defined as the age when the first derivative became positive. **E**, Ages at the start of developmental increase across three connection types: sensorimotor-sensorimotor (SS), sensorimotor-association (SA), and association-association (AA). These categories refer respectively to connections within sensorimotor networks, between sensorimotor and association networks, and between association networks. Start ages were significantly delayed in ADHD relative to TD for SA (T = 3.9, *P* < 0.001) and AA (T = 5.8, *P* < 0.001) connections. **F**, Connection-wise lead/lag in developmental timing for the ADHD group compared with the TD group. Light red indicates a developmental delay, and light blue indicates a developmental advance. White indicates that start ages for both groups fell outside the observed window (before 8.9 years or after 15.5 years), so the difference could not be inferred. ^n.s^: not significant; SC: structural connectivity; ADHD: attention-deficit/hyperactivity disorder; TD: typically developing; S: sensorimotor; A: association.

We next estimated the age at which each connection began to increase in strength (**Fig. 2D**). Consistent with hierarchical connectivity development^12^, TD showed earlier maturation of sensorimotor connections and later maturation of association connections. Critically, ADHD participants exhibited a further delay in this onset for association-involving connections. We categorized the connections into three types: sensorimotor-sensorimotor (SS), sensorimotor-association (SA), and association-association (AA). These three categories refer respectively to connections within sensorimotor networks, between sensorimotor and association networks, and between association networks. Ages at the start of SC-strength increase were significantly delayed in ADHD for both SA (*T* = 3.9, *P* < 0.001) and AA (*T* = 5.8, *P* < 0.001) connections (**Fig. 2E**). We then compared start ages between ADHD and TD groups for each individual connection (**Fig. 2F**) and found that delays were concentrated in connections linking higher-order association networks. By contrast, most within sensorimotor connections began increasing before the lower age bound of our sample (8.9 years) for both groups, which limits interpretation of developmental lead or lag for these connections within the observed window. These results demonstrated that SC development shows a lag in the maturation of the higher-order association connectivity.

### Structural connectivity deviations align with the S-A connectional axis

Having demonstrated delayed maturation of SC in ADHD relative to TD youth, we next quantified the extent to which individual’s connectivity deviated from the TD-derived normative developmental trajectories (**Fig. 3A**). For each participant and visit, observed SC strength was compared with the age-expected distribution defined by the normative trajectories from the TD training set to obtain its centile, which were then transformed into a Z score, yielding deviation scores for both the TD test and ADHD groups. Averaging across participants yielded group-level deviation maps. Youths with ADHD exhibited more positive deviations, particularly in connections with association networks (**Fig. 3B**), whereas the TD test group showed minimal deviation from the normative reference (**Fig. 3C**).

**Fig. 3.**
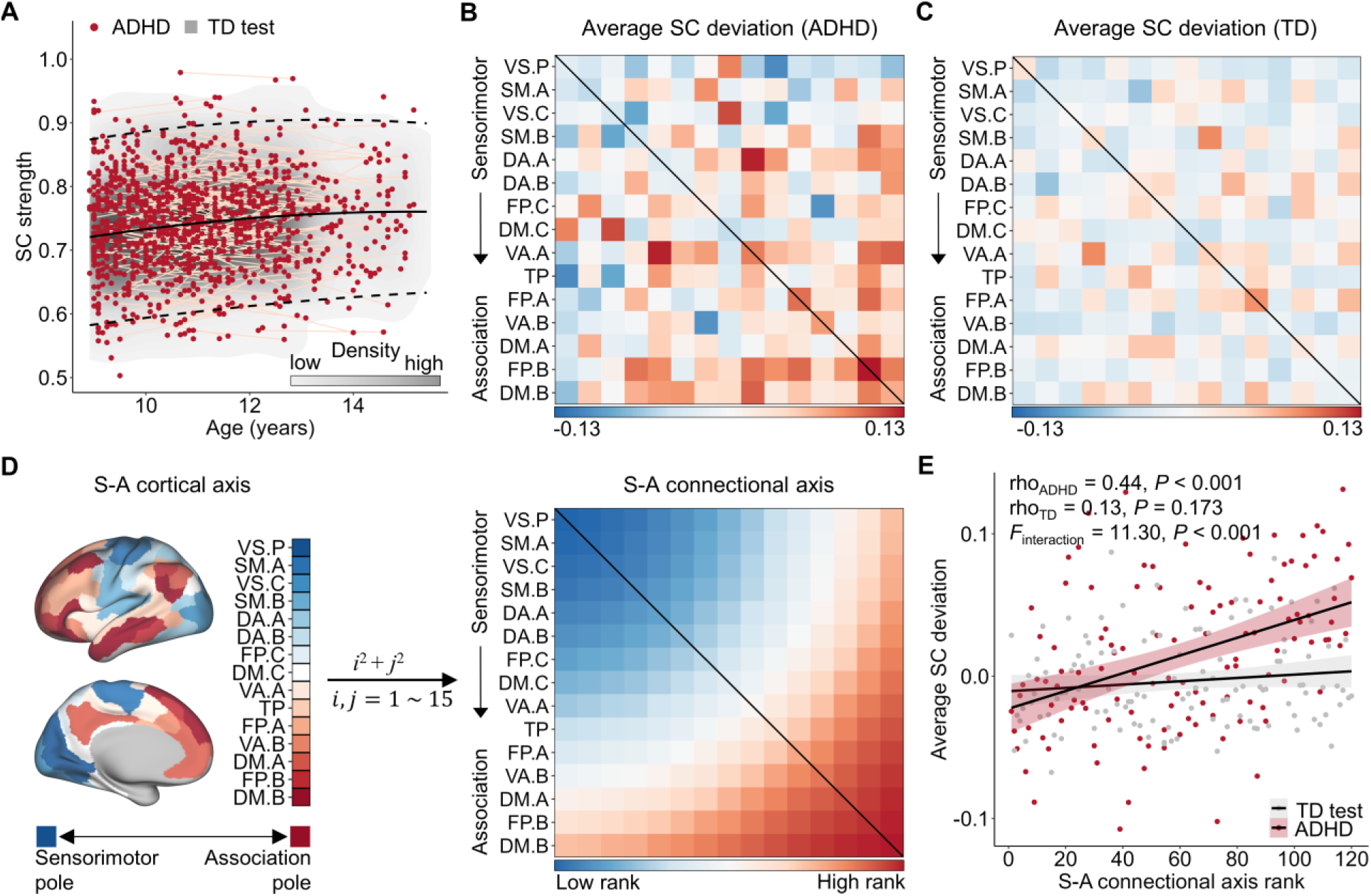
Structural connectivity deviations are organized along the sensorimotor-association connectional axis. **A**, Schematic of the normative modeling procedure using the mean SC strength across the connectome as an example. The solid black line represents the age-expected median trajectory derived from the TD training set, with dashed lines indicating the 2.5^th^ and 97.5^th^ percentiles. Red points (ADHD) and a gray density (TD test) show observed SC strength relative to the normative range. Deviation scores were calculated to quantify how far each observation diverged from the age-expected distribution predicted by the normative model. **B-C**, Group-averaged deviation matrices across the 15 cortical networks for ADHD (**B**) and TD (**C**). In ADHD, positive deviations indicated stronger-than-expected connectivity, particularly within higher-order association connections. **D,** The S-A connectional axis was derived from the canonical S-A cortical axis. Each of the 15 cortical networks was assigned a median S-A rank based on its constituent vertices. A single connectional axis rank was then computed for each connection by summing the squared ranks of its paired networks and rescaled to a discrete range of 1–120. **E**, Mean SC deviations showed a significant positive correlation with S-A connectional axis ranks in ADHD (Spearman’s rho = 0.44, *P* < 0.001) but not in TD youths (rho = 0.13, *P* = 0.173). An interaction analysis confirmed a significant group difference in alignment with the S-A connectional axis (*F* = 11.30, *P* < 0.001). SC: structural connectivity; S-A: sensorimotor-association; ADHD: attention-deficit/hyperactivity disorder; TD: typically developing.

To examine the spatial organization of these deviations, we mapped them along the canonical S-A axis^20^, a hierarchical gradient extending from unimodal, primary sensorimotor to transmodal, higher-order association cortices. Each of the 15 cortical networks was assigned a rank according to its position along the cortical S-A axis (**Fig. 1C** left; **Fig. 3D** left). Following our previous approach^25^, we then derived an S-A connectional axis by summing the squared cortical ranks of the two networks linked by each connection and rescaling the resulting values to a 1-120 range (**Fig. 3D**). This framework situates connections along a continuous gradient, with lower ranks corresponding to early-developing sensorimotor connections and higher ranks to late-maturing association connections that support complex integrative functions.

In youths with ADHD, group-mean SC deviations showed a significant positive correlation with S-A connectional axis ranks (Spearman’s rho = 0.44, *P* < 0.001), indicating that larger deviations occurred in higher-order association connections positioned toward the transmodal end of the axis (**Fig. 3E**). No such relationship was observed in the TD test group (rho = 0.13, *P* = 0.173). An interaction analysis further confirmed that this alignment with the S-A connectional axis was significantly stronger in ADHD (*F* = 11.30, *P* < 0.001). Together, these findings demonstrate that SC alterations in ADHD are not uniformly distributed across the connectome but are preferentially concentrated within higher-order association connections that support integrative cognitive functions.

### Developmental decline of structural connectivity deviations in ADHD

Symptoms of ADHD often decline during adolescence^2,3,26^, raising the possibility that atypical brain connectivity may also evolve with age. Having established delayed maturation of SC strength in ADHD relative to TD youth, we next examined whether these deviations from normative development changed across youth within the ADHD group. For each connection, we fitted a generalized additive mixed model (GAMM) with SC deviation as the dependent variable, including a thin-plate spline for age and random intercepts for participants, while controlling for sex and head motion. The partial *R*²of the smooth age term quantified the effect size of developmental change for each connection. To capture the direction of change, we assigned the sign to the partial *R*²based the average first derivative of the fitted trajectory, with negative values indicating age-related decreases in deviation and positive values indicating increases (**Fig. 4A**).

**Fig. 4.**
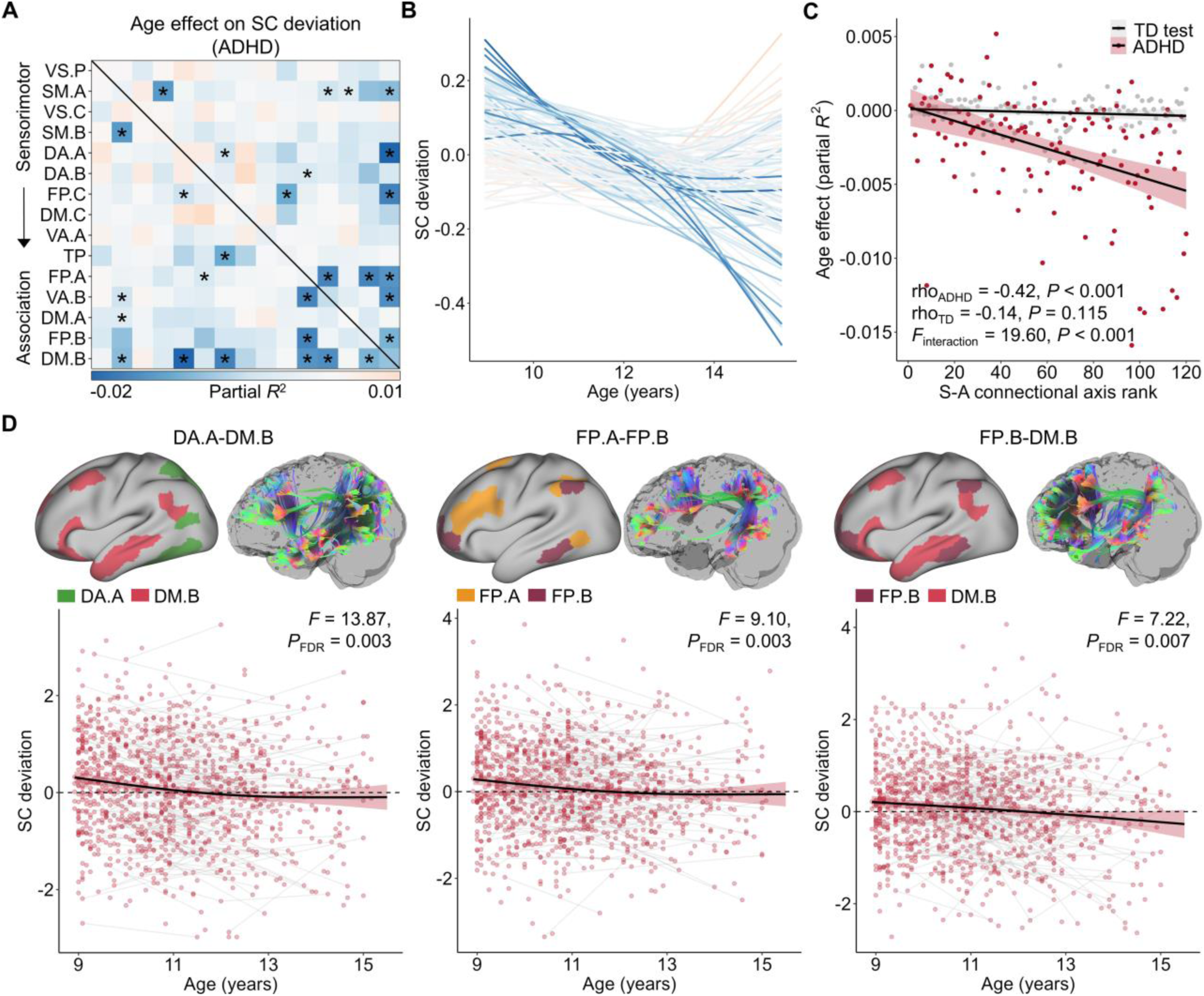
Age-related changes of SC deviations in ADHD. **A**, Age effects (partial *R*^2^) on SC deviations across 120 connections in youths with ADHD. Fourteen connections showing significant ADHD-related age effects (*P*_FDR_ < 0.05) are marked with black asterisks. A connection was considered ADHD-related if it showed a significant age effect within the ADHD group and a significantly different age-by-diagnosis interaction compared with the TD test group. **B**, Developmental trajectories of SC deviations in ADHD. Each curve represents one connection, color-coded by its corresponding effect size from panel (A). **C**, The magnitude of age effects on SC deviations showed a significant alignment with the S-A connectional axis in ADHD (Spearman’s rho = -0.42, *P* < 0.001), but not in TD youths (rho = -0.14, *P* = 0.115). An interaction analysis confirmed that this alignment with the S-A axis was significantly stronger in ADHD (*F* = 19.60, *P* < 0.001). **D**, Example trajectories for three representative connections (DA.A-DM.B, FP.A-FP.B, and FP.B-DM.B) randomly selected from the 14 significant ones in panel (A). SC deviations decreased with age, shifting from positive values in childhood toward values not significantly different from zero by mid-adolescence, indicating age-related normalization of connectivity. DA-A: Dorsal attention-A; FP-A: Frontoparietal-A; FP-B: Frontoparietal-B; DM-B: Default mode-B. SC: structural connectivity; S-A: sensorimotor-association; ADHD: attention-deficit/hyperactivity disorder; TD: typically developing.

Across the connectome, most connections in the ADHD group exhibited decreasing SC deviations with age (**Fig. 4B**). Twenty-five connections showed significant age-related decreases in deviation (*P*_FDR_ < 0.05). To determine which of these effects were related to ADHD, we tested age-by-diagnosis interaction, identifying 14 connections that were both significant within ADHD and showed stronger age effects than in TD (*P*_FDR, interaction_ < 0.05; **Fig. 4A**, **Table S3**). These ADHD-related connections were largely concentrated in higher-order association networks, including frontoparietal, dorsal-attention, and default-mode networks. We next assessed whether this developmental attenuation followed the S-A connectional axis. The signed age-effect sizes (partial *R*²) were significantly correlated with S-A connectional axis rank in ADHD (Spearman’s rho = −0.42, *P* < 0.001), indicating that the strongest age-related reductions in deviation occurred in connections positioned toward the association end of the axis (**Fig. 4C**). No such correlation was found in TD youths (rho = -0.14, *P* = 0.115). A diagnoses-by-rank interaction confirmed that this alignment with the S-A axis was significantly stronger in ADHD (*F* = 19.6, *P* < 0.001).

Representative trajectories for three connections between dorsal attention, frontoparietal, and default mode networks showed SC deviations declining steadily from childhood through mid-adolescence, approaching the normative range (their 95% CIs did not cross zero; **Fig. 4D**). Notably, although these examples remain above zero, several other connections (6 out of 14) exhibited deviations that crossed zero over the age span. This pattern suggests that atypical connectivity in ADHD diminishes with age but does not fully normalize, consistent with prior evidence that most individuals retain residual symptoms into adulthood. Collectively, these findings indicate that SC alterations in ADHD are developmentally dynamic, showing partial convergence toward typical patterns yet persistent reorganization in higher-order association connections that subserve integrative cognitive control.

### SC deviations mediate and track symptom improvement in ADHD

Having established that SC deviations decline with age in ADHD, we next investigated whether these developmental changes relate to symptom trajectories. ADHD-related behavioral symptoms were assessed using parent-reported T scores from the Child Behavior Checklist (CBCL) ^27,28^, focusing on two dimensions: ADHD problems, reflecting overall symptom burden, and attention problems, capturing executive attentional control difficulties. These two dimensions derive from the DSM-oriented and syndrome scales, respectively, and together index overall ADHD symptom burden. Developmental changes in both dimensions were modeled using GAMMs with sex as a covariate. Consistent with prior epidemiological findings^26^, both ADHD problems (*F* = 4.31, *P*_FDR_ = 0.015, **Fig. 5A**) and attention problems (*F* = 8.79, *P*_FDR_ < 0.001, **Fig. 5B**) showed significant age-related decreases across adolescence.

**Fig. 5.**
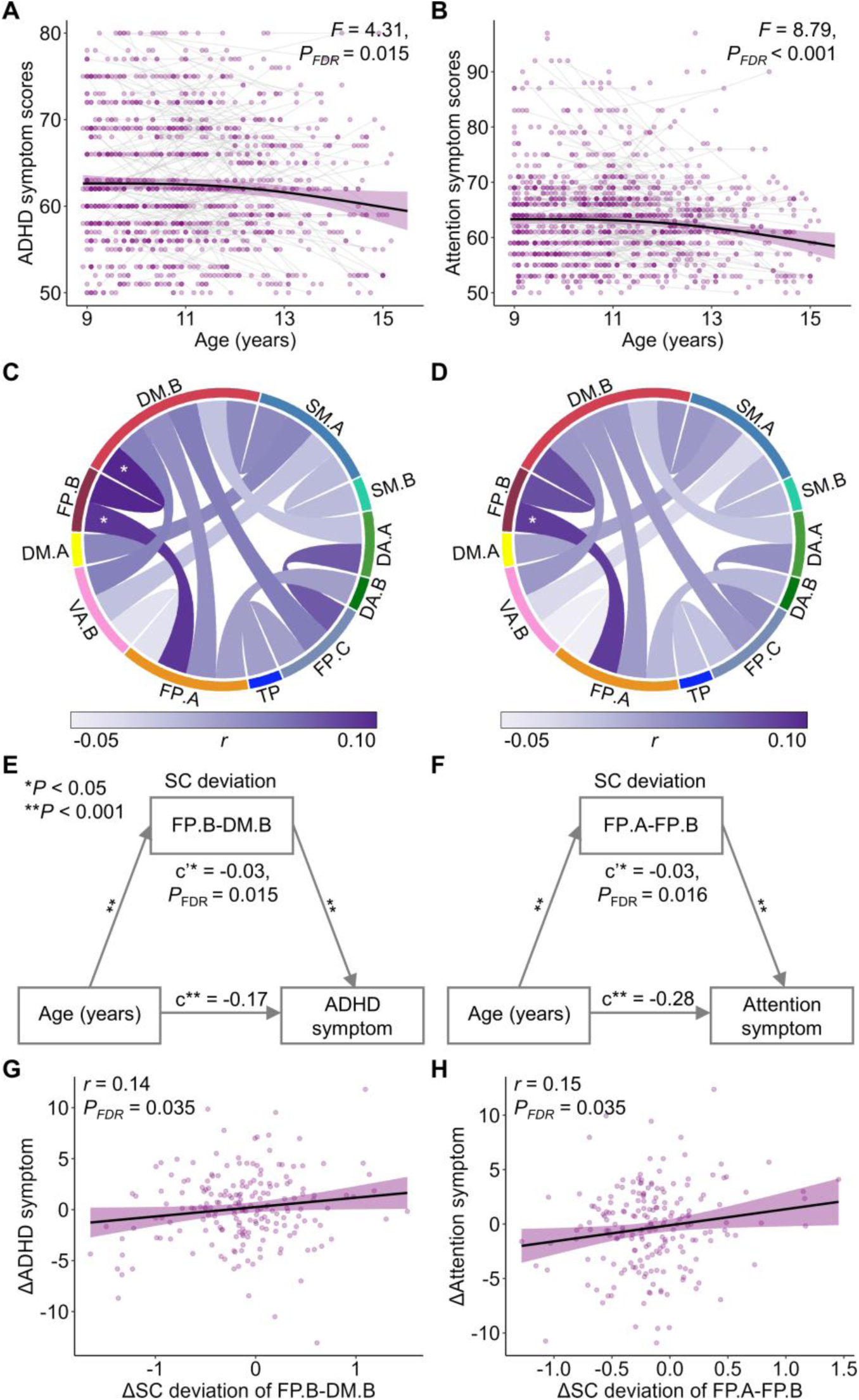
SC deviations mediate and track symptom improvement in ADHD. **A**–**B**, Both ADHD problems (**A**) and attention problems (**B**) decreased significantly with age in youths with ADHD (*P*_FDR_ < 0.05). Parent-reported T scores were derived from the Child Behavior Checklist (CBCL), where ADHD problems reflect overall clinical symptom burden and attention problems reflect executive attentional control difficulties. **C–D**, Circos plots show brain-symptom associations across the 14 connections that exhibited ADHD-related developmental decline in SC deviation (Fig. 4A). Significant positive associations (*P*_FDR_ < 0.05) are highlighted by asterisks: SC deviations in FP.A–FP.B and FP.B–DM.B correlated with ADHD problems (**C**), and deviations in FP.B–DM.B correlated with attention problems (**D**). **E–F**, SC deviations of FP.B–DM.B mediated the age-related decline of ADHD problems (**E**), and SC deviations of FP.A–FP.B mediated the age-related decline of attention problems (**F**). Path coefficients (a, b, c, c’) are shown with significance levels indicated (*: *P* < 0.05, **: *P* < 0.001). **G–H**, Within-subject analyses showing that reductions in SC deviations were correlated with parallel reductions in symptoms: FP.B–DM.B with ADHD problems (**G**) and FP.A–FP.B with attention problems (**H**). FP.A: Frontoparietal control A; FP.B: Frontoparietal control B; DM.B: Default mode B. SC: structural connectivity; ADHD: attention-deficit/hyperactivity disorder.

We then evaluated whether age-related reductions in SC deviations were related to these behavioral improvements. Focusing on the 14 connections that showed ADHD-related age effects (**Fig. 4A**), we fit GAMMs with SC deviation as a predictor of symptom scores, including a smooth term for age, sex, and head motion as covariates, and participant as a random intercept. Significant positive associations emerged between SC deviations in FP.A-FP.B (*r* = 0.09, *P*_FDR_ = 0.036) and FP.B-DM.B (*r* = 0.10, *P*_FDR_ = 0.036) connections and ADHD problems (**Fig. 5C**), and between SC deviation in FP.B-DM.B connection and attention problems (*r* = 0.09, *P*_FDR_ = 0.044, **Fig. 5D**), indicating that greater deviations were associated with more severe symptoms.

Next, we used bootstrapped mediation analyses to test whether SC deviations mediated the age-related decline in symptoms while adjusting for sex and head motion. Deviations in the FP.B-DM.B connection significantly mediated reductions in ADHD symptom (indirect effect = -0.03, *P_FDR_* = 0.015; **Fig. 5E**), whereas deviations in the FP.A-FP.B connection mediated reductions in attention problems (indirect effect = -0.03, *P_FDR_* = 0.016; **Fig. 5F**). To validate these mediation effects at the individual level, we tested whether within-person changes in SC deviations tracked longitudinal changes in symptoms. For participants with two or three visits (N = 176) in the ADHD group, developmental rates were computed for SC deviations and symptom scores as the difference between visits divided by the inter-visit interval, yielding 215 observations. Within-subject analyses confirmed that reductions in SC deviations were associated with decreases in symptom severity. Specifically, changes in SC deviation for FP.B-DM.B connection correlated with changes in ADHD problems (*r* = 0.14, *P*_FDR_ = 0.035; **Fig. 5G**), and changes in SC deviation for FP.A-FP.B connection correlated with changes in attention problems (*r* = 0.15, *P*_FDR_ = 0.035; **Fig. 5H**). In sum, our results demonstrated that SC deviations within frontoparietal–default mode circuitry partially mediated the developmental improvement of ADHD symptom.

### Developmental delays of SC deviations are reproducible across cohorts

We next tested whether the developmental and hierarchical patterns of SC deviations observed in the ABCD cohort generalized to an independent Chinese cohort comprising 292 TD youths and 468 youths with ADHD (ages 6.5-15.5 years; **Fig. 1B**). Consistent with the discovery sample, the onset ages of SC strengthening were delayed in ADHD compared to TD for both sensorimotor-association (SA; *T* = 2.3, *P* < 0.05) and association-association (AA; *T* = 2.2, *P* < 0.05) connections (**Fig. 6A**). We next examined whether the spatial organization of SC deviations along the S-A connectional axis was reproduced. In ADHD, the group-mean deviation of each connection, averaged across participants, showed a significant positive correlation with its S-A rank across all 120 connections (Spearman’s rho = 0.19, *P* = 0.036, **Fig. 6B**), indicating that connections closer to the association end of the axis exhibited larger deviations.

**Fig. 6.**
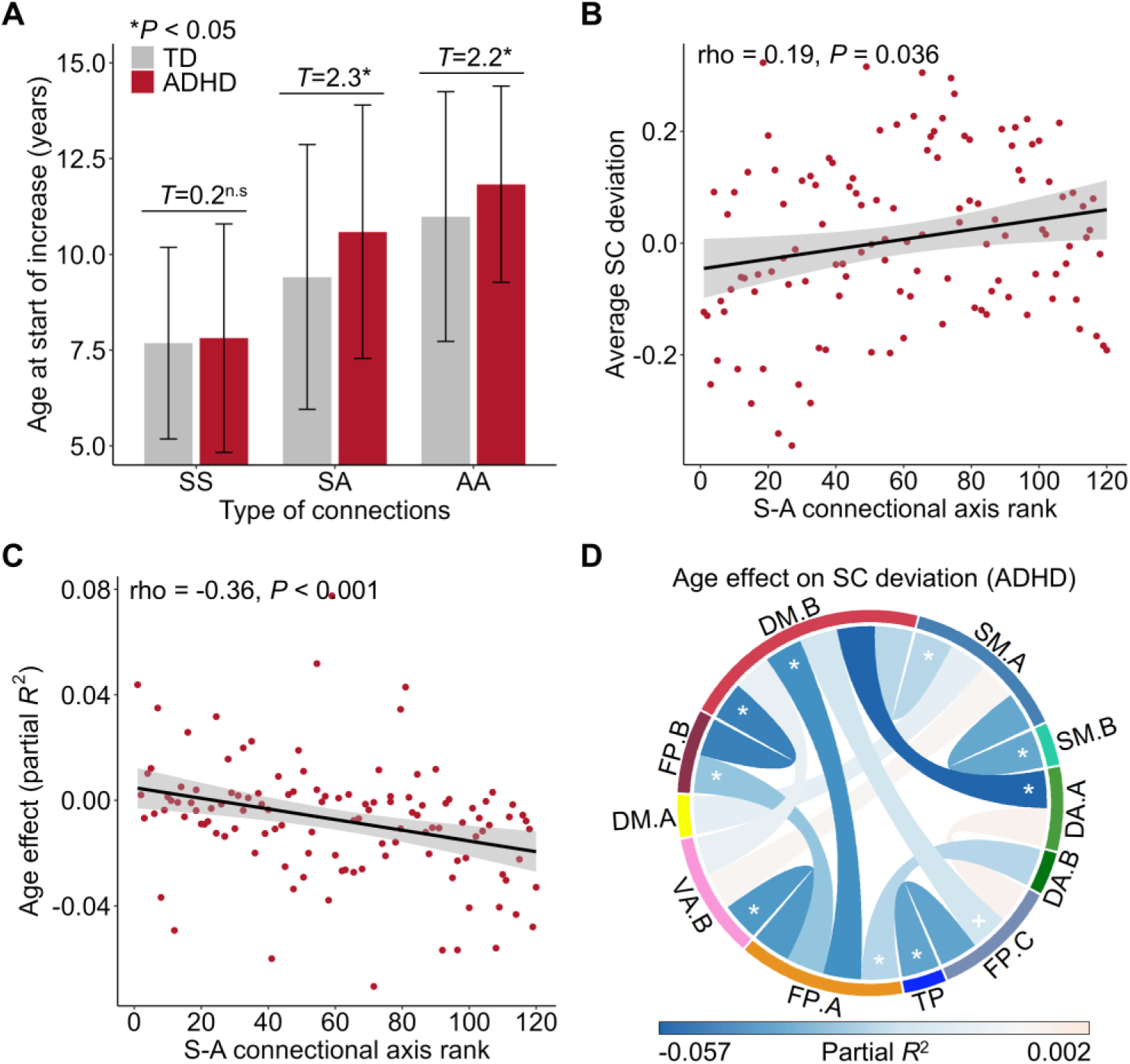
Developmental delays of SC deviations were replicated in an independent Chinese cohort. **A,** Ages at the start of SC strengthening across three connection types, including sensorimotor-sensorimotor (SS), sensorimotor-association (SA), and association-association (AA), in youths with ADHD and TD controls. Consistent with the ABCD discovery cohort, start ages of developmental increase were significantly delayed in ADHD relative to TD for SA (*T* = 2.3, *P* = 0.031) and AA (*T* = 2.2, *P* = 0.023) connections. **B**, Group-average SC deviations were significantly correlated with S-A connectional axis ranks across 120 connections (rho = 0.18, *P* = 0.036), indicating larger deviations in higher-order association connections. **C**, The magnitudes of age effects on SC deviations showed a significant negative correlation with the S-A connectional axis ranks (rho = -0.36, *P* < 0.001). **D**, Of the 14 connections that showed ADHD-related decreases in deviation in the ABCD study, 10 also exhibited significant age-related reductions in deviation in the Chinese cohort. ^n.s^: not significant; *: *P*_FDR_ < 0.05, ^+^: *P*_FDR_ < 0.10. SC: structural connectivity; S-A: sensorimotor-association; ADHD: attention-deficit/hyperactivity disorder.

We then examined the reproducibility of developmental age effects on SC deviations. For each connection, we estimated the effect size (partial *R*^2^) of developmental changes of connectivity strength and signed it based on the average first derivative. Across the connectome, these age effects were significantly negatively correlated with S-A connectional axis ranks (rho = -0.36, *P* < 0.001, **Fig. 6C**), indicating that the strongest developmental reductions in SC deviations occurred in higher-order association connections. We next focused on the 14 connections that showed ADHD-related age effects in the ABCD cohort (**Fig. 4A**) to assess their reproducibility. Of these, nine exhibited significant age-related reductions in deviations in the Chinese cohort (*P_FDR_* < 0.05), for example DA.A-DM.B, FP.B-DM.B, FP.A-DM.B connections, and one exhibited a marginally significant age effect (FP.C-DM.B, *P_FDR_* = 0.056; **Fig.6D**, **Table S4**).

Harmonized CBCL symptom measures were unavailable in the Chinese cohort, precluding direct replication of symptom associations. Nonetheless, the consistent developmental delays and hierarchical organization of SC deviations across two large and culturally distinct populations demonstrate the robustness and generalizability of the observed neurodevelopmental patterns.

### SC deviations predict and normalize with pharmacological treatment

Having established that developmental normalization of SC deviations parallels symptom improvement in ADHD, we next investigated their clinical relevance in a pharmacological treatment cohort from the Peking University Sixth Hospital (PKU6). This cohort included 112 youths with ADHD (ages 7.0-15.5 years) who completed dMRI, T1WI, and ADHD rating scale (ADHD-RS) assessments before treatment (**Fig. 7A**). The ADHD-RS is a validated parent-reported questionnaire comprising 18 items corresponding to DSM-5 criteria for inattention and hyperactivity-impulsivity, with higher scores indicating greater symptom burden^29^. Among these participants, 72 received methylphenidate (MPH) and 40 received atomoxetine (ATX). After 12 weeks of medication, all participants completed post-treatment ADHD-RS evaluations, and 37 underwent follow-up dMRI and T1WI scans (**Fig. 7A**). Both medications produced significant symptom reductions, with inattention and hyperactivity-impulsivity scores decreasing after treatment (all *P* < 0.001; **Fig. 7B**) and comparable improvement across drug groups (inattention: *F* < 0.001, *P* = 0.981; hyperactivity-impulsivity: *F* = 1.99, *P* = 0.161).

**Fig. 7.**
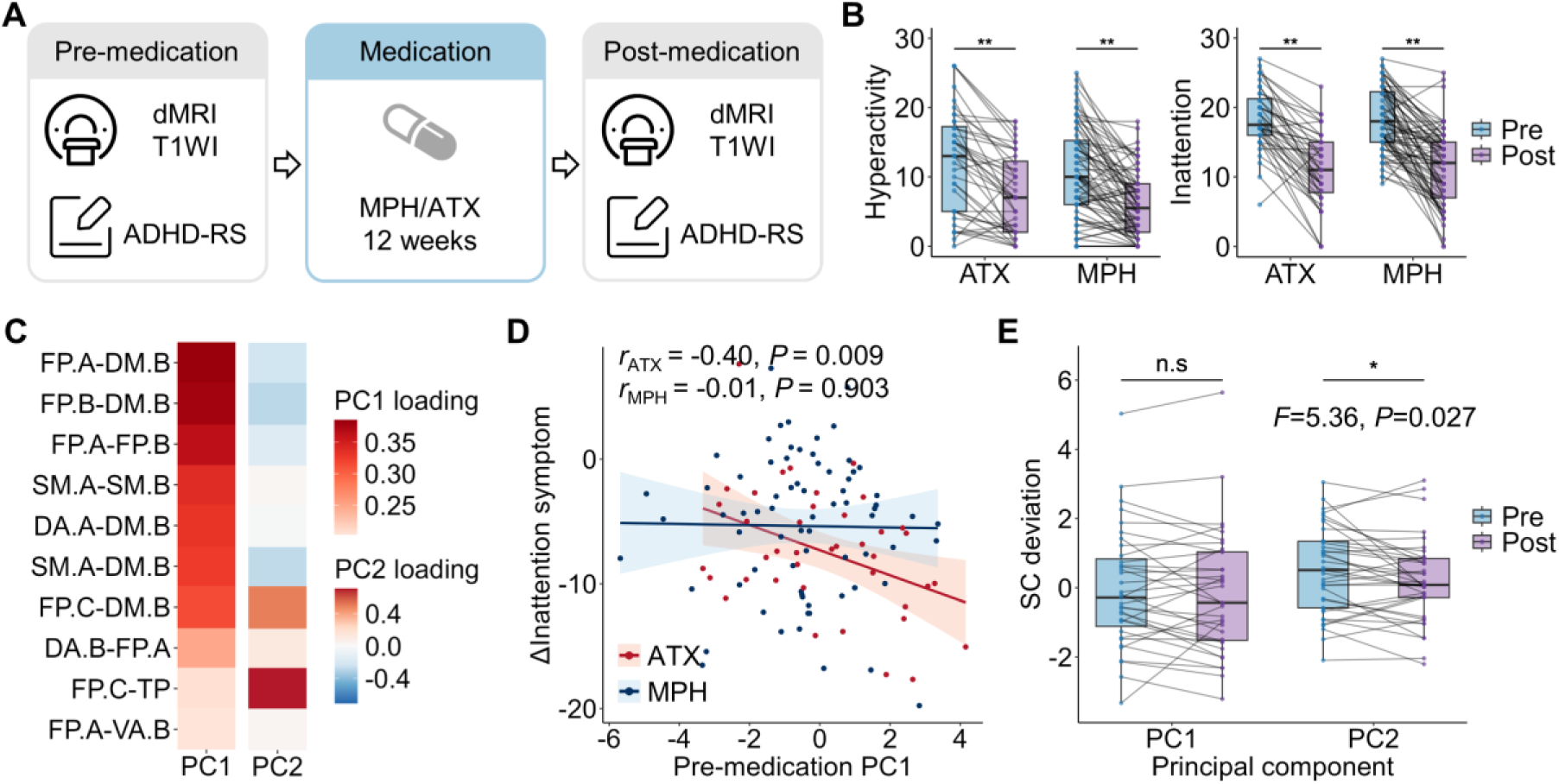
Associations between SC deviations and medication treatment response. **A**, Schematic of the medication study protocol in the PKU6 cohort. A total of 112 children with ADHD completed dMRI, T1WI, and ADHD-RS assessments before treatment. Of these, 72 received MPH and 40 received ATX. After 12 weeks of medication, all participants completed the ADHD-RS, and 37 underwent follow-up dMRI and T1WI scans. **B**, Both ATX and MPH treatments significantly reduced hyperactivity and inattention symptoms (*P* < 0.001 for both). **C**, Principal component analysis (PCA) of SC deviations across the 10 replicated connections (from Fig. 6D). The heatmap shows component loadings for the first two principal components (PC1 and PC2). **D**, Baseline PC1 scores significantly predicted greater reductions in inattention symptoms following ATX treatment (*r* = -0.40, *P* = 0.009, *P*_FDR_ = 0.037), but not MPH (*r* = -0.01, *P* = 0.903, *P*_FDR_ = 0.903). **E**, Across both drug groups combined (n = 37), repeated-measures ANOVA revealed a significant post-treatment reduction in PC2 scores (*F* = 5.36, *P* = 0.027, *P*_FDR_ = 0.053), indicating decreased SC deviations following medication. PKU6: Peking University Sixth Hospital; dMRI: diffusion MRI; T1WI: T1-weighted imaging; ADHD-RS: ADHD-rating scales; MPH: methylphenidate; ATX: atomoxetine; SC: structural connectivity; PC: principal component.

We then tested whether baseline SC deviations predicted 12-week symptom improvement, defined as the change in ADHD-RS scores from pre- to post-treatment. Focusing on the 10 connections that showed reproducible developmental effects in the PKU6 cohort (**Fig. 6D**), we applied principal component analysis (PCA) to summarize individual baseline deviation profiles. The first two components (PC1 and PC2) explained 45.6% and 13.0% of the variance, respectively (**Fig. S5**). PC1 primarily reflected SC deviations in frontoparietal**–**default mode connectivity (FP.A-DM.B, FP.A-FP.B, FP.B-DM.B), whereas PC2 captured frontoparietal**–**temporal-parietal and frontoparietal–default mode connectivity (FP.C-TP, FP.C-DM.B). Baseline PC1 scores significantly predicted subsequent improvement in inattention symptoms among children treated with ATX (*r* = -0.40, *P* = 0.009), but not among those treated with MPH (*r* = -0.01, *P* = 0.903; **Fig. 7D**). Greater baseline deviations were associated with stronger inattention improvement for atomoxetine. No significant relationship was observed for hyperactivity-impulsivity symptom.

To test whether pharmacotherapy altered SC deviation over time, we examined the 37 participants with longitudinal imaging data. A repeated-measures ANOVA revealed a significant post-treatment reduction in PC2 deviation scores (*F* = 5.36, *P* = 0.027; **Fig 7E**), indicating partial normalization of connectivity deviations following medication. Because of the limited sample size, participants from both drug groups were analyzed together. No significant effect was observed for PC1. Together, these results indicate that distinct components derived from connections showing ADHD-related developmental reductions differentially relate to pharmacological outcomes: PC1 predicts response to atomoxetine, whereas PC2 reflects treatment-induced normalization of SC.

## Discussion

This study provides large-scale, longitudinal, and cross-cultural evidence that developmental deviations in white matter SC characterize youths with ADHD. Across two independent cohorts, ADHD was marked by deviations from normative trajectories of cortico-cortical connectivity, particularly in higher-order association pathways linking frontoparietal, default-mode, and attention networks. These deviations diminished through adolescence, paralleling the decline in ADHD symptoms, and longitudinal reductions in connectivity deviations tracked individual differences in symptom improvement. Baseline SC deviations predicted treatment response to atomoxetine but not methylphenidate, and medication treatment further normalized the connectivity deviations. Together, these findings identify SC deviations as potential developmental markers of ADHD with translational relevance for personalized intervention.

Our results extend prior evidence of delayed cortical maturation in ADHD^13,14^ by demonstrating that similar developmental lags exist at the level of white-matter connectivity. The preferential involvement of higher-order association pathways aligns with the hierarchical organization of brain development^12,20^, in which sensorimotor circuits mature early while association systems remain plastic into adolescence. This protracted plasticity may render association pathways particularly vulnerable to atypical development while also retaining potential for adaptive reorganization. The observed decline in connectivity deviations with age indicates that white matter abnormalities in ADHD are not static but evolve across development, reflecting a delayed or altered maturational trajectory rather than a fixed neurodevelopmental deficit. Importantly, the connections showing the strongest normalization primarily localized in frontoparietal and default-mode networks, which are critical for cognitive control and internal mentation, domains often impaired in ADHD^9^.

The magnitude of SC deviations linking the frontoparietal and default mode networks mediated the developmental decline of ADHD symptoms and tracked within-subject symptom change over time. These large-scale association networks support inhibitory control, sustained attention, and other higher-order executive functions^30,31^, that are commonly impaired in ADHD^32^. Our symptom–connectivity associations showed that greater deviations in frontoparietal and frontoparietal–default mode pathways correspond to more severe symptoms, providing white-matter evidence that disruptions in these networks contribute to the clinical phenotype of ADHD. Prior longitudinal research shows that a substantial proportion of children with ADHD exhibit partial symptomatic remission during adolescence^2,33^. Our within-individual analyses suggest a potential neural explanation: youths who show greater normalization of connectivity deviations in these association pathways tend to experience greater symptom improvement, whereas those whose deviations persist or worsen show more stable or worsening clinical trajectories. These results identify frontoparietal–default mode pathways as central to symptom-related developmental change in ADHD.

Connectivity deviations are also linked clinical change. In the Chinese cohort, baseline deviations of cortical SC predicted treatment response to atomoxetine, but not methylphenidate. This dissociation aligns with their distinct neuropharmacological profiles: atomoxetine, a selective norepinephrine reuptake inhibitor, preferentially enhances cortical catecholaminergic signaling^34^, whereas methylphenidate engages subcortical dopaminergic circuits more strongly^35,36^. The predictive dimension of SC deviations (PC1), centered on frontoparietal–default mode circuits, identified youths who later showed greater symptom reduction with atomoxetine, suggesting that atypicality within these higher-order networks indexes a cortical substrate of treatment susceptibility. In contrast, a sperate dimension of deviations (PC2) showed significant reductions following 12 weeks of medication, indicating that pharmacological intervention can partially normalize SC during adolescence. This dissociation suggest that distinct components of SC deviations capture complementary aspects of pathophysiology, with PC1 reflecting a trait-like marker of sensitivity to cortical norepinephrine modulation, and PC2 reflecting a more state-dependent substrate that exhibits short-term neuroplastic responses to treatment. These findings are consistent with evidence that monoaminergic signaling influences myelination and axonal remodeling^37^. Together, these results highlight SC deviations as promising markers for stratifying treatment and monitoring therapeutic effects.

Methodologically, this work advances beyond prior diffusion MRI studies of ADHD that have yielded heterogeneous results^15^. Traditional case–control designs compare static group averages and ignore normative developmental variation, while tract-based analyses lack specificity regarding the cortical regions they connect, limiting interpretability. By applying a connectome-wide normative modeling framework, we quantified person-specific deviations in system-level connectivity trajectories across development. The large sample size, longitudinal design, and replication in an independent, cross-cultural cohort enhance the robustness and generalizability of the findings. These results underscore the value of integrating normative modeling with developmental and pharmacological data to identify clinically meaningful neural markers in neuropsychiatric disorders.

Several limitations warrant consideration. First, accurately reconstructing individual white-matter SC from diffusion MRI remains technically challenging. We mitigated this limitation by applying state-of-the-art probabilistic fiber tractography using multi-shell, multi-tissue constrained spherical deconvolution^38^, combined with anatomically constrained tractography^39^ and spherical deconvolution-informed filtering of tractograms^40^ to improve biological accuracy. Second, while our cohorts spanned childhood through mid-adolescence, extending this approach into early adulthood will be crucial to determine whether connectivity deviations fully normalize or persist in individuals with enduring symptoms. Finally, although we focused on ADHD as a single diagnostic entity, the disorder is clinically heterogeneous and often comorbid with other neurodevelopmental conditions; future work incorporating transdiagnostic dimensions may better capture shared and distinct developmental pathways.

In conclusion, this study demonstrates that ADHD involves developmental deviations in large-scale white-matter connectivity that parallel symptom trajectories and predict pharmacological outcomes. These findings bridge neurodevelopmental and clinical perspectives on ADHD, highlighting adolescence as a critical window for structural brain maturation and therapeutic intervention. By integrating normative modeling with longitudinal and cross-cultural data, this work established a scalable framework for mapping individual developmental profiles and advancing precision psychiatry in neurodevelopmental disorders.

## Methods

### Participants

This study leveraged data from the Adolescent Brain Cognitive Development (ABCD) study as the discovery dataset and an independent Chinese cohort, which served both as a validation sample and for the medication analyses. The ABCD study is a multi-site, longitudinal project that recruited approximately 10,000 children aged 9–10 years across the United States at baseline^18^. We accessed neuroimaging data from the ABCD Fast-Track Portal in July 2024 and demographic and clinical information from ABCD Release 5.1. Based on the parent-reported computerized version of the Kiddie Schedule for Affective Disorders and Schizophrenia (KSADS-COMP)^41^ for DSM-5, we included typically developing (TD) children with no diagnoses and children diagnosed with attention-deficit/hyperactivity disorder (ADHD)^42^. Only participants with complete MRI data were included. Initially, we included 7,731 scans from TD children (*N*_baseline_=3,743, *N*_2year-follow-up_=3,259, *N*_4year-follow-up_=731) and 1,454 scans from children with ADHD (*N*_baseline_=808, *N*_2year-follow-up_=517, *N*_4year-follow-up_=129). Then, we applied the following exclusion criteria: 1) lack of self-fluency in English or lack of parental fluency in English or Spanish; 2) severe sensory, intellectual, medical, or neurological conditions; 3) missing age or sex information; 4) comorbid with schizophrenia, major depressive disorder or substance abuse; 5) not meeting the official imaging inclusion criteria outlined in Release 5.1 (see **Supplementary Text** for details); 6) failure in data processing; 7) excessive head motion (mean framewise displacement (FD) > Mean+3×SD); 8) whole-brain structural connectivity (SC) strength outside the normal range (Mean±3×SD) within each group; 9) sites with a sample size of fewer than 30 scans in the TD group. After screening, we finally included 6,687 scans from TD children aged 8.9 to 15.5 years (3,323 males, *N*_baseline_=3,155, *N*_2year-follow-up_=2,878, *N*_4year-follow-up_=654) and 1,114 scans from children with ADHD aged 8.9 to 15.2 years (800 males, *N*_baseline_=597, *N*_2year-follow-up_=401, *N*_4year-follow-up_=116). The study protocol was approved by the institutional review board of the University of California, San Diego. Legal guardians provided written informed consent, and children gave verbal assent.

The Chinese cohort combined data from two studies: the Executive Function and Neurodevelopment in Youth (EFNY) study and a study from Peking University Sixth Hospital (PKU6)^43^. Both studies recruited TD children and children with ADHD. ADHD was diagnosed using the parent-reported Kiddie-Schedule for Affective Disorders & Schizophrenia, Present & Lifetime Version (K-SADS-PL)^44^. Inclusion criteria for both studies were: 1) normal intelligence (EFNY: Raven’s IQ score > 80, PKU6: full-scale IQ score > 80, assessed with the Wechsler Child Intelligence Scale, Third Edition); 2) no MRI contraindications; 3) no comorbid axis I disorder other than tic disorder or oppositional defiant disorder; and 4) no severe physical or neurological conditions; 5) no history of psychotic medication use. We restricted the sample to participants aged 6.5–15.5 years due to the small number of participants outside this age range. Initially, 306 TD children (*N*_EFNY_=205, *N*_PKU6_=101) and 481 children with ADHD (*N*_EFNY_=38, *N*_PKU6_=443) with complete MRI data were included. Participants with hearing problems or excessive head motion and whole-brain SC strength outside the normal range were subsequently excluded, resulting in a final sample of 292 TD children aged 6.5 to 15.4 years (*N*_EFNY_=195, *N*_PKU6_=97) and 468 children with ADHD aged 6.5 to 15.5 years (*N*_EFNY_=38, *N*_PKU6_=430). Ethical approval for the EFNY study was obtained from the Human Research Ethics Committee of the Chinese Institute for Brain Research, Beijing, and the protocol for the PKU6 study was approved by the Ethics Committee of Peking University Sixth Hospital.

The inclusion and exclusion flow charts are presented in **Fig. S1,S2**, the age distributions are presented in **Fig. 1A,B**, and demographic details are available in **Table S1,S2**.

### Symptom assessment

We assessed clinical symptoms of children with ADHD using the parent-reported school-age Child Behavior Checklist (CBCL/6-18) from the ABCD study. The CBCL consists of 113 items evaluating youth emotional and behavioral problems and demonstrates strong reliability and validity^27,28^. It provided eight domain-specific syndrome scales, such as anxious/depressed, somatic complaints, and attention problems. In 2003, six DSM-Oriented scales (e.g., ADHD problems, affective problems) were developed based on expert clinical consensus^45,46^. In this study, we selected the attention scale and the ADHD scale to evaluate the symptoms of children with ADHD. To examine age-related changes in clinical symptoms while controlling for typical developmental effects, we used CBCL T-scores, with higher T-scores indicating more severe symptoms relative to age- and sex-specific norms.

The PKU6 study from the Chinese cohort leveraged parent-reported ADHD-rating scales (ADHD-RS)^29^ to evaluate the clinical symptoms of children with ADHD before and after the medication treatment. ADHD-RS contains 18 items corresponding to the DSM-5 criteria for ADHD, which can be divided into two subscales: inattention and hyperactivity/impulsivity. The higher scores indicate more severe symptoms.

### MRI acquisition

This study used T1-weighted imaging (T1WI) and diffusion MRI (dMRI) data from each participant at each visit. ABCD data were acquired across multiple sites on 3.0T scanners from Siemens, GE, and Philips^47^. EFNY data were acquired on 3.0T Siemens scanners, while data from the PKU6 study were acquired on a GE Discovery MR750 3.0T scanner. Detailed acquisition parameters are summarized in **Table S5**.

### MRI data processing and structural connectome reconstruction

We applied the preprocessing pipelines embedded in QSIPrep version 0.16.0RC3 (https://qsiprep.readthedocs.io/)^48^ to the T1WI and dMRI data. QSIPrep configures standardized pipelines for processing both diffusion and structural MRI data, incorporating algorithms from *Nipype* 1.8.1^49^, *ANTs* 2.3.1^50^, *FSL* 6.0.5^51^, *MRtrix3*^52^, *Nilearn* 0.9.1^53^, and *Dipy*^54^. For the T1WI data, preprocessing included: 1) intensity non-uniformity correction; 2) skull stripping; 3) spatial normalization. Then, we reconstructed pial surface and segmented tissues from the intensity-normalized T1WI data using *FreeSurfer* 7.1.1^55^, which provided anatomical constraints for tractography.

For the dMRI data from the ABCD and EFNY studies, we first concatenated the dMRI and the b0 image with reversed phase-encoding direction. This step was not applied to the PKU6 study because reverse phase-encoded b0 images were not collected. The dMRI data then underwent the following preprocessing steps: 1) designating frames with a b-value less than 100 s/mm^2^ as b = 0 volumes; 2) Marchenko-Pastur principal component analysis (MP-PCA) denoising^56^; 3) Gibbs unringing^57^; 4) B1 bias field correction^58^; 5) head motion, susceptibility distortion, and eddy current corrections^59^; 6) coregistration to individual T1WI and realignment to the ACPC orientation. Distortion correction was not applied to the PKU6 data due to the absence of reverse phase-encoded b0 images. Following previous studies^60^, mean FD was computed as the sum of the absolute values of the differentiated realignment estimates for each volume to quantify head motion during the dMRI scans.

After preprocessing, we performed whole-brain probabilistic fiber tracking on each dMRI scan using MRtrix3^52^. Because the dMRI data from the ABCD and EFNY studies were acquired with multi-shell parameters, we applied multi-shell multi-tissue constrained spherical deconvolution (CSD)^38,61^ to estimate the fiber orientation distribution (FOD) in each voxel. For the PKU6 data, which were acquired with single-shell parameters, we applied single-shell 3-tissue CSD to estimate voxel-wise FODs^61^. Afterwards, 1) FOD images were intensity normalized; 2) Whole-brain tractography was conducted under the anatomically constrained tractography (ACT) framework based on the hybrid surface and volume segmentation (HSVS) algorithm^39,62^. The ACT framework, which leverages *FreeSurfer*-derived pial surfaces and tissue segmentations, enhances the biological accuracy of tractography. Tractography was performed using the iFOD2 algorithm^63^ with four FOD samples per step, generating 10 million streamlines ranging from 30 mm to 250 mm in length. To match streamline densities with fiber densities estimated by CSD, we applied spherical-deconvolution informed filtering of tractograms (SIFT2) to assign an appropriate cross-sectional area multiplier to each streamline^40^. SC matrices were then constructed based on the Yeo-17 network^64^, merged from the Schaefer-400 atlas^23^. Because the limbic networks typically exhibit low signal-to-noise ratios^19^, they were excluded, resulting in 15 remaining networks. A radial search (maximum radius = 2 mm) was performed from each streamline endpoint to identify the nearest gray matter node. To reduce false-positive streamlines in global tractography^65^, we applied a consistency-based thresholding method^66^ on the region-level connectome. Streamlines belonging to connections with a coefficient of variation (CV) above the 75^th^ percentile were removed^67,68^. Finally, for each participant at each visit, we constructed a 15×15 large-scale network with 120 undirected connections. The weight of each connection, defined as structural connectivity (SC) strength, was calculated as the number of streamlines divided by the arithmetic mean volume of the connected node pair^24^.

### Normative modeling

Following the recommendations of the World Health Organization, we used generalized additive models for location, scale and shape (GAMLSS)^21,22^ to estimate the normative models for SC strength of each connection. This procedure was completed via *gamlss* package^69^ in R (version 4.2.2). Compared with traditional generalized models, GAMLSS can model not only the mean (*μ*) but also the scale (*σ*) and shape (*v*, *τ*) for the response variable. These normative models were then used to estimate standardized SC deviation scores (z scores) for clinical individuals (ADHD group) at each visit. In line with a standard protocol^70^, it is important to retain a proportion of healthy control participants (TD group) in the test set. Accordingly, we split the TD participants from the ABCD study into training (70%, N=4,599, 2,285 males; N_baseline_ = 2,177; N_2-year follow-up_ = 1,984; N_4-year follow-up_ = 438) and test (30%, N=2,088, 1,038 males; N_baseline_ = 978; N_2-year follow-up_ = 894; N_4-year follow-up_ = 216) datasets, stratified by sex, site, and longitudinal data availability. However, due to the limited sample size of the TD group (N=292) in the Chinese cohort, all TD participants were used to train the models. All sites in the training set included > 30 observations. The following GAMLSS estimation steps were conducted using the TD training set. We also fit normative models in all children with ADHD using the same parameters as in the TD set to compare developmental features between groups.

#### (1) Selection of the optimal distribution and spline parameters

From over 100 distributions available in the GAMLSS framework, we evaluated 25 continuous distribution families with three or four parameters (*μ*, *σ*, *v*, *τ*), as listed in **Table S6**. In our models, age was included as a smoothing term, modeled with B-spline function, while sex and head motion (mean FD) were included as fixed effects. A random effect for site was also included. These covariates were used to model the location (*μ*) and scale (*σ*) parameters, while only intercepts were fitted for the shape parameters (*v*, *τ*). The optimal distribution for each edge was determined by selecting the model with the lowest Bayesian Information Criterion (BIC) that successfully converged. The distribution most frequently selected across all connections was then chosen as the final optimal distribution for the entire dataset. For the ABCD dataset, the generalized gamma (GG) distribution was selected. For the Chinese cohort, the Box–Cox Power Exponential (BCPEo) distribution, with a log link for the *μ* parameter, was chosen (**Fig. S3A**). All models were estimated using maximum likelihood, with convergence defined as a change in log-likelihood of less than 0.001 across successive iterations (up to a maximum of 200).

Recognizing that age effects on brain phenotypes are often nonlinear, we modeled the effect of age on the location (*μ*) and scale (*σ*) parameters using B-spline basis functions. The complexity of a B-spline is determined by its degrees of freedom (df) and the degree of its piecewise polynomial. To identify the optimal parameters, we fitted GAMLSS models with a range of df from 2 to 5 and a range of polynomial degrees from 2 to 3. The optimal parameter set was identified based on the lowest BIC and successful model convergence. The set most frequently selected across all connections was defined as the final optimal parameter set. For both datasets, df=2 and polynomial degree=2 were selected (**Fig. S3B**).

#### (2) Normative model fitting

For each connection in the 15×15 large-scale network, we modeled the SC strength distributions via GAMLSS. Based on prior analysis, we selected the GG distribution for the ABCD dataset and the Box-Cox Power Exponential (BCPEo) distribution for the Chinese cohort dataset, using the default link function for each parameter. The final model for each connection was defined as follows:

ABCD:

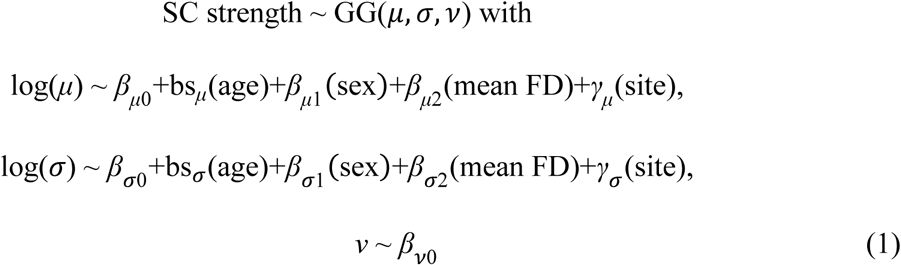

Chinese cohort:

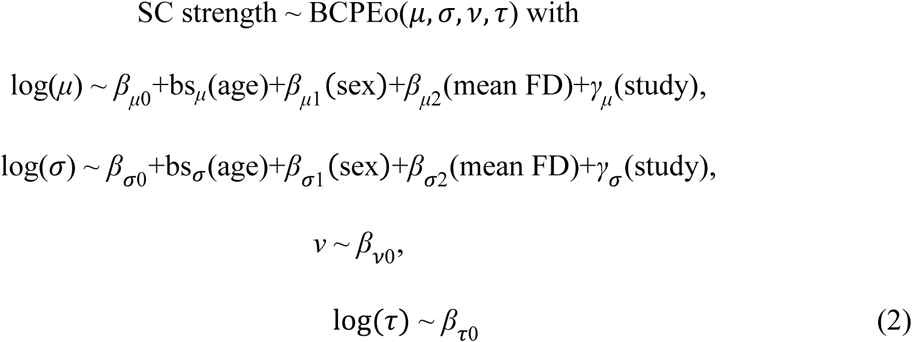

For each connection, the SC strength served as the response variable. The model included age, sex, and mean FD as fixed effects, with site or study included as a random effect. Based on prior analysis, age was modeled as a smoothing term using a B-spline basis function with two degrees of freedom and a second-degree polynomial.

To characterize developmental features, we derived two summary metrics for each connection’s trajectory: the average first derivative and the age at start of increase. To quantify overall age effects on SC strength, we estimated the first derivative of the median developmental trajectory at 1,000 evenly spaced ages across the observed range using finite differences^71^, then averaged these derivatives over the full range. We defined the age at start of increase as the age at which the first derivative crossed zero from negative to positive. For between-group comparisons, we classified connections into three types: SS (sensorimotor-sensorimotor; VS.P, VS.C, SM.A, SM.B), SA (sensorimotor-association), and AA (association-association; all remaining systems). We compared ages at start of increases between TD and ADHD using paired-wise T-tests within each connection type. Boundary handling followed consistent rules. If a trajectory was already increasing at the lower bound of the dataset, we set the start age to that minimum (ABCD: 8.9 years; Chinese cohort: 6.5 years). If a trajectory did not begin increasing within the observed range, we set the start age to the maximum (15.5 years for both datasets).

#### (3) Model evaluation

To evaluate the goodness-of-fit for our GAMLSS models, we assessed the distribution of the normalized quantile residuals^22,72^. First, we generated Q-Q (quantile-quantile) plots and Owen’s plots for the model of mean SC strength across the entire network (ABCD: **Fig. S4A,B**; Chinese cohort: **Fig. S4E,F**). For both the ABCD and Chinese cohort datasets, the Q-Q plots of the normative quantile residuals demonstrated that the sample quantiles closely followed the theoretical quantiles, indicating that the residuals were approximately normally distributed. In the Owen’s plots, the 95% confidence intervals consistently encompassed the horizontal zero line, suggesting no systematic deviation from the expected distribution. Next, we computed the skewness and kurtosis of the normative quantile residuals for the model of each individual connection. For both datasets, the skewness coefficients for all models were within the range of - 0.5 to 0.5, and the excess kurtosis coefficients were within the range of 2 to 4 (ABCD: **Fig. S4C**; Chinese cohort: **Fig. S4G**), further supporting the approximate normality of the residuals and the adequacy of the model assumptions.

To assess the models’ reliability, we performed a bootstrap analysis for each connection model. This involved 1,000 bootstrap iterations using stratified sampling with replacement. The resampling was stratified by site (or study, for the Chinese cohort) and sex to maintain the proportional representation of the original datasets. For each iteration, a GAMLSS model was fitted with the same parameters as the primary analysis to derive a median normative trajectory across the age range. To assess the convergence of the model fits across all bootstrap iterations, we computed the coefficients of variation (CV) at 1,000 equally-spaced age points and then averaged them. For all connection models in both datasets, the average CV did not exceed 0.1 (ABCD: **Fig. S4D**; Chinese cohort: **Fig. S4H**), indicating good model reliability.

#### (4) Individual deviation quantification

We computed individual deviations in SC strength for both TD participants in the test set and ADHD patients using the GAMLSS models fitted to the TD training set. For each connection, we first extracted individual centile scores from the age-, sex-, and mean-FD–specific normative distributions^73,74^. These centile scores were then transformed into deviation scores using randomized quantile residuals. For convenience, we refer to these as SC deviation scores. Each participant’s SC deviation score reflects how extreme an individual’s SC strength is relative to the reference distribution.

### Definition of the S-A connectional axis

The S-A cortical axis provided a hierarchical framework of human brain, spanning continuously from primary sensorimotor to transmodal association cortices^20^. In this study, we sought to characterize the spatial distribution of SC deviations in ADHD patients and how these deviations change with age within the framework of the S-A axis. We first ranked the Yeo-17 networks according to the median S-A cortical axis rank of the vertices within each network (**Fig. 1C**). Then, building on our previous work^25,75^, we extended the S-A cortical axis into the connectional dimension. As shown in equation (3), we defined the element (*C_i,j_*) in the matrix of the S-A connectional axis ranks as the quadratic sum of the S-A cortical axis ranks of *node_i_* and *node_j_*. We then computed the ranks of *C* to obtain the S-A connectional axis ranks.

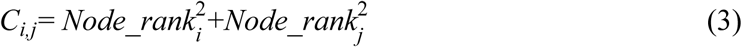

In the context of a 15×15 large-scale connectome, the S-A connectional axis rank ranges from 1 to 120 (**Fig. 3D**).

### Statistical analysis

#### (1) Alignment of SC deviations with the S-A connectional axis

First, we computed the mean SC deviation across observations for each connection within the 15×15 Yeo network, performing this calculation separately for the ADHD group and the TD test group. We then used Spearman’s rank correlation to assess the alignment between this mean SC deviation map and the S-A connectional axis map. To formally test whether this alignment was specific to the ADHD group, we fitted a linear regression model that included an interaction term between the S-A connectional axis ranks and diagnostic group for the ABCD dataset. A significant interaction term would indicate that the relationship between SC deviation and the S-A ranks is moderated by diagnosis, thus confirming specificity to children with ADHD. For the Chinese cohort, we did not perform the interaction analysis because a separate TD test dataset was not created due to the small sample size.

#### (2) Development of deviations in SC strength

We examined age-related changes in SC deviations among children with ADHD across the developmental period from middle childhood to middle adolescence. Given the longitudinal design of the ABCD dataset, we applied generalized additive mixed models (GAMMs) using the *gamm4* package^76^ in R4.2.2. In each model, SC deviation was the response variable, and nonlinear effects of age were modeled while controlling for sex, mean FD, and subject-specific random intercepts. Smooth plate regression splines were used for the smooth term, with smoothing parameters estimated via restricted maximum likelihood. Following prior neurodevelopmental studies, the spline basis dimension (*k*) was set to 3^77,78^. The model for each connection was specified as:

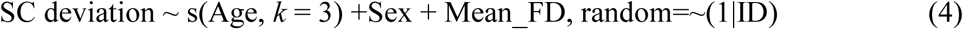

The significance of age effects was evaluated by comparing the full model with a null model (excluding age)^78,79^ using parametric bootstrap likelihood ratio tests with 10,000 simulations (via the *pbkrtest* package^80^). Resulting *P* values were adjusted for multiple comparisons using the false discovery rate (FDR), with a significance threshold of 0.05. Effect sizes of age were quantified using partial *R*^2^ between the full and null models. Finally, we tested whether the spatial distribution of age effects aligned with the S-A connectional axis using Spearman’s correlation. Consistent with the analyses above, we further examined the interaction between the S-A ranks and diagnosis using a linear regression model.

For connections showing significant age-related changes in SC deviation, we further tested whether these developmental effects were specific to participants with ADHD by modeling the interaction between age and diagnosis (formula 5). A combined dataset of children with ADHD and TD participants from the test set was used.

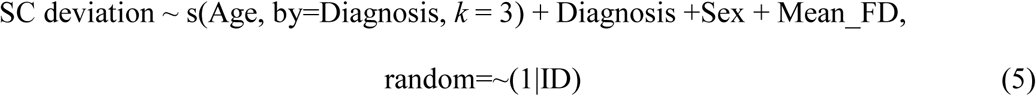

The significance of the interaction term was assessed by comparing the full model with a null model (excluding the interaction) using parametric bootstrap likelihood ratio tests with 10,000 simulations^79^ and resulting *P* values were corrected using the FDR method. For connections with significant age effects detected in the ABCD dataset, we validated the results in the Chinese cohort using generalized additive models (GAM) with similar model specifications. Comparisons with null models were performed using parametric bootstrap testing via analysis of variance (ANOVA, implemented in the *ecostats* package^81^).

### Mediation analysis of the ADHD-related symptoms

Following the developmental analysis, we conducted a mediation analysis to investigate whether SC deviations mediate the development of ADHD-related symptoms, as measured by the CBCL. This analysis was conducted only in children with ADHD. We first tested the effect of age on each of the two CBCL symptom scores using a GAMM, with sex included as a covariate. Model significance was determined by comparison to a null model that excluded the age term. *P*-values were corrected for multiple comparisons across the three symptoms. To establish the necessary preconditions for mediation, we further tested the association between symptoms and the SC deviations of connections that showed significant age effects. GAMM was used for this analysis, controlling for age, sex, and mean FD, as shown in formula (6). Significance was determined by comparing this model to a null model that excluded the SC deviation term. *P*-values were corrected using the FDR method, and partial correlation coefficients were computed to quantify the effect size.

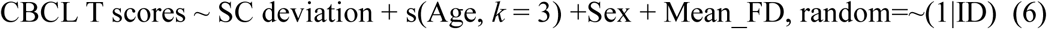

Connections with SC deviations that were significantly associated with symptoms were selected as potential mediators in a multiple-mediator model, implemented using the *lavaan* package^82^. Because the *lavaan* package cannot directly incorporate random effects, we first regressed out the subject-specific random intercepts from the data using linear mixed-effects models from the *lme4* package. The residuals (*M*) from these models were then used in the subsequent mediation analyses. In addition, for each symptom category, the independent variable was age (*X*), and the dependent variable was the symptom score (*Y*). The significance of the indirect effects was examined using bootstrapped confidence intervals for 10,000 iterations. To identify the most parsimonious set of mediators, we first fitted a full mediation model. We then fitted a reduced model containing only the mediators that were significant in the full model. The full and reduced models were compared using a likelihood-ratio test (ANOVA). If the reduced model significantly outperformed the full model, it was selected for the final analysis. Finally, the *P*-values for the indirect effects of each mediator in the final models were corrected across the two symptom categories using the FDR method.

To further validate the mediation effects, we conducted a within-subject analysis including only participants with longitudinal measurements (N=176, 138 males) in the ADHD group. Developmental rates were calculated as the change in SC deviation or symptom scores divided by the interval between visits, for example, (SC deviation_Time2_ - SC deviation_Time1_)/(Age_Time2_ - Age_Time1_). Rates were computed for each visit pair (2-year vs. baseline, 4-year vs. baseline, and 4-year vs. 2-year), yielding 215 data points. Multiple linear regression was then used to examine the relationship between symptom developmental rates and SC deviation rates, controlling for average age, sex, and average mean FD. Significance of SC deviations was assessed by comparing full models with null models using ANOVA. *P*-values were corrected for multiple comparisons across the two symptoms, and partial correlation coefficients were calculated to quantify effect sizes.

### Associations between SC deviations and treatment response

We further examined the association between SC deviations and medication response in children with ADHD. This analysis focused on the connections showing ADHD-specific, age-related changes in SC deviations to assess the clinical relevance of our findings. In the PKU6 study, 112 children with ADHD (86 males, 7.0–15.5 years) underwent 12 weeks of medication treatment with either methylphenidate (MPH, N = 72) or atomoxetine (ATX, N = 40) following the baseline MRI scan. Treatment commenced the morning after baseline, with an initial dosage of 18 mg/day for MPH and 10 mg/day for ATX. Dosages were adjusted based on clinical response and adverse effects—weekly for MPH and every 2–4 days for ATX (maximum dosage: 54 mg/day for MPH; 1.2 mg/kg or 100 mg/day for ATX). All treatments were prescribed and monitored by a professional child psychiatrist (Q.J.C. or L.Y.). ADHD symptoms were assessed before and after treatment using the ADHD-RS. Of the 112 children, 37 also underwent a follow-up MRI scan after taking their usual morning medication. The study protocol was presented in **Fig. 6A** (clinical trial ID: NCT05229627).

We tested whether baseline SC deviations could predict treatment response. A principal component analysis (PCA) was performed on SC deviations across different connections in children with ADHD from the Chinese cohort to extract metrics reflecting overall variance of SC deviations. The first two components (PC1 and PC2), explaining 58.53% of the variance, were selected as general SC deviation metrics. Linear regression models were used to examine whether baseline SC deviation PC1 and PC2 predicted symptom changes (post-treatment − pre-treatment ADHD-RS scores), with analyses conducted separately for the MPH and ATX groups. The significance was evaluated by ANOVA analysis with null models, which only contained covariates. We next evaluated whether SC deviation PC1 and PC2 changed following medication treatment using repeated-measures ANOVA. We tested for main effects of time, medication type, and their interaction. Finally, we examined associations between changes in SC deviation principal components (post-treatment − pre-treatment) and symptom improvements using linear regression models. Partial correlation coefficients were calculated to quantify effect sizes.

## Data and code availability

The ABCD dataset used in this study is available to qualified researchers upon application through the ABCD Data Repository (see https://abcdstudy.org/scientists/data-sharing/). The Chinese cohort data used in this study are not currently available, as data collection for both projects is still ongoing. The EFNY dataset will be made publicly available within one year. All analysis scripts are available at https://github.com/CuiLabCIBR/ADHDdeviation.git. All analysis methods are described in the main text and supplementary materials.

## Acknowledgements

We thank the research participants and staff involved in data collection and project executive of the Adolescent Brain Cognitive Development (ABCD) Study, the Executive Function and Neurodevelopmental in Youth (EFNY), and the Peking University Sixth Hospital study (PKU6). The ABCD data were provided by the ABCD Study (abcdstudy.org) funded by NIH. The ABCD Study is a multisite, longitudinal study designed to recruit more than 10,000 children ages 9-10 years old and follow them over 10 years into early adulthood. The ABCD Study is supported by the NID and additional federal partners under award numbers U01DA041048, U01DA050989, U01DA051016, U01DA041022, U01DA051018, U01DA051037, U01DA050987, U01DA041174, U01DA041106, U01DA041117, U01DA041028, U01DA041134, U01DA050988, U01DA051039, U01DA041156, U01DA041025, U01DA041120, U01DA051038, U01DA041148, U01DA041093, U01DA041089, U24DA041123 and U24DA041147. A full list of supporters is available at https://abcdstudy.org/federal-partners.html. A listing of participating sites and a complete listing of the study investigators can be found at https://abcdstudy.org/consortium_members/. ABCD consortium investigators designed and implemented the study and/or provided data but did not necessarily participate in the analysis or writing of this report. This manuscript reflects the views of the authors and may not reflect the opinions or views of the NIH or ABCD consortium investigators. The ABCD data repository grows and changes over time. The EFNY study were conducted by our lab in Chinese Institute for Brain Research, Beijing, supported by the STI 2030-Major Projects (grant No.2022ZD0211300). The PKU6 study was assisted by the National Center for Protein Sciences at Peking University for assistance with MRI experiments in the MRI center of Peking University Sixth hospital.

## Author contributions

Z.C. and X.X. conceptualized the study. X.X. conducted the formal analysis. Z.F. designed the medication study. L.Y, Q.C., Z.F., K.Z., Y.Z. and X.Z. acquired PKU6 dataset. Z.C., Y.L., X.X., and H.X. acquired EFNY dataset. H.X. replicated all analyses. Z.C. managed the project administration and supervised the project. H.Y., A.K., T.X., R.C., W.C., Q.C., and L.Y. provided comments. X.X. and Z.C. wrote the original draft. All authors reviewed and edited the manuscript.

