## Supplementary Materials for "Developmental deviations of structural connectivity in youths with ADHD predict symptom and treatment outcomes"

**Supplementary information**

Supplementary Text

Fig. S1 to Fig. S5

Table S1 to S6

**Supplementary Text**

ABCD official imaging inclusion criteria for dMRI:

A participant's scan was included if all of the following were satisfied: 1) at least one diffusion magnetic resonance imaging (dMRI) series was complete and passed quality control; 2) the complete dMRI series contained 103 volumes, or for Philips two-run acquisitions, 51 volumes per run; 3) at least one T1-weighted (T1WI) series was complete and passed quality control; 4) B0 field maps for distortion correction were available; 5) FreeSurfer quality control was not failed; 6) manual post-processing quality control for dMRI was not failed; 7) dMRI successfully registered to T1WI; 8) field of view was complete (no truncation); 9) derived dMRI outputs were available.

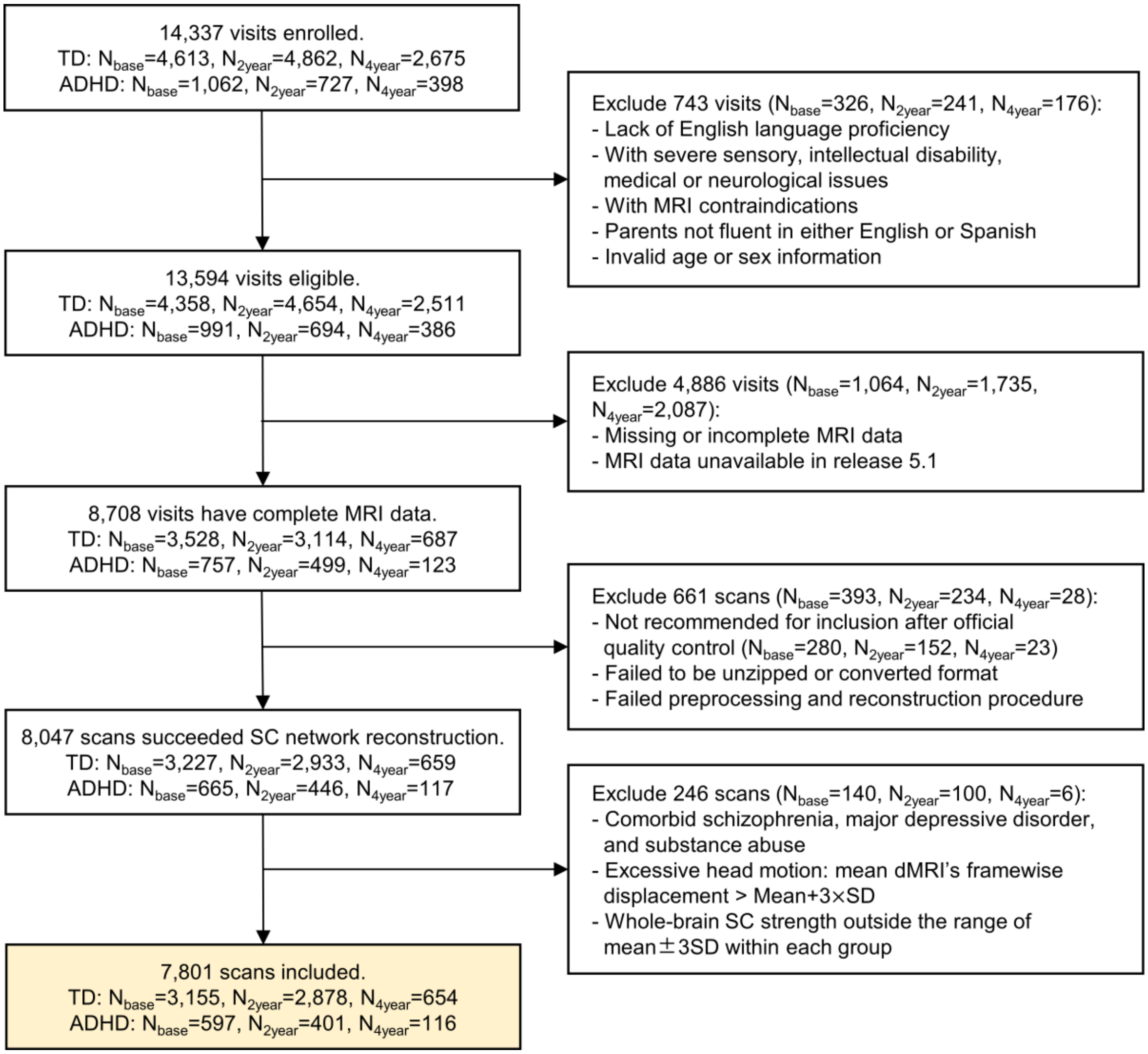


**Fig. S1 | Flowchart of inclusion and exclusion for participants in the ABCD dataset.** ABCD: Adolescent Brain Cognitive Development; ADHD: attention-deficit/hyperactivity disorder; TD: typically developing; SD: standard deviation.

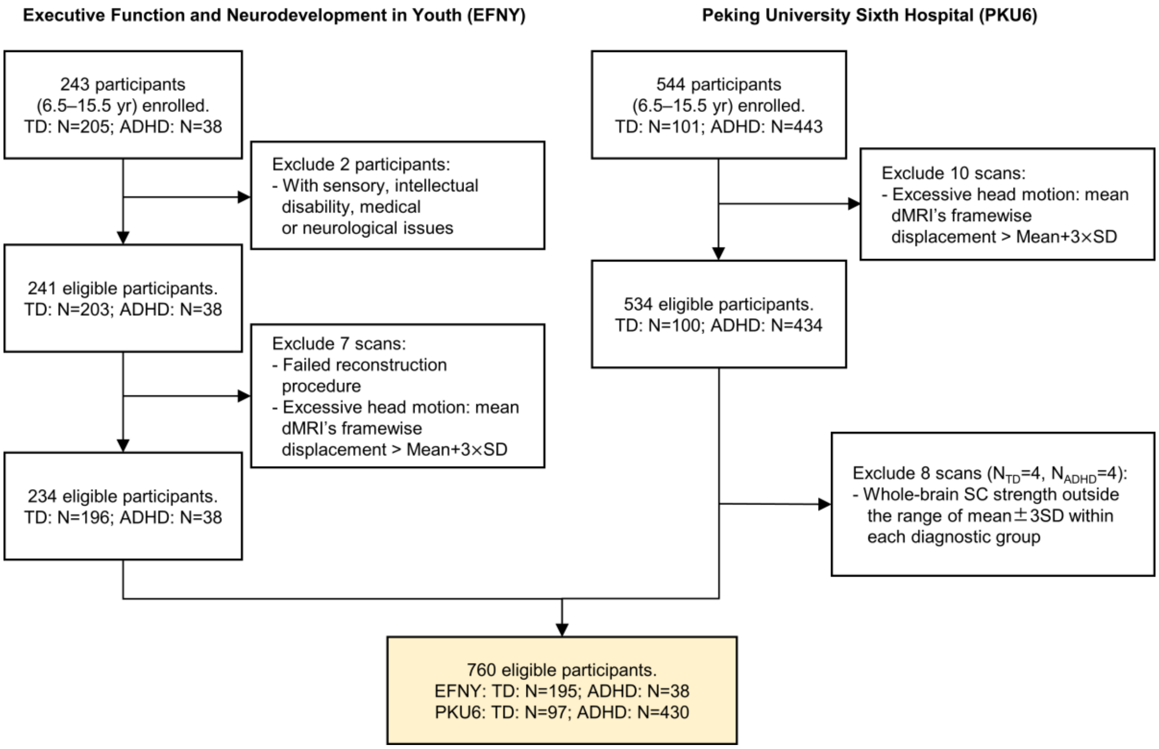


**Fig. S2 | Flowchart of inclusion and exclusion for participants in the Chinese Cohort.** ADHD: attention-deficit/hyperactivity disorder; TD: typically developing; SD: standard deviation.

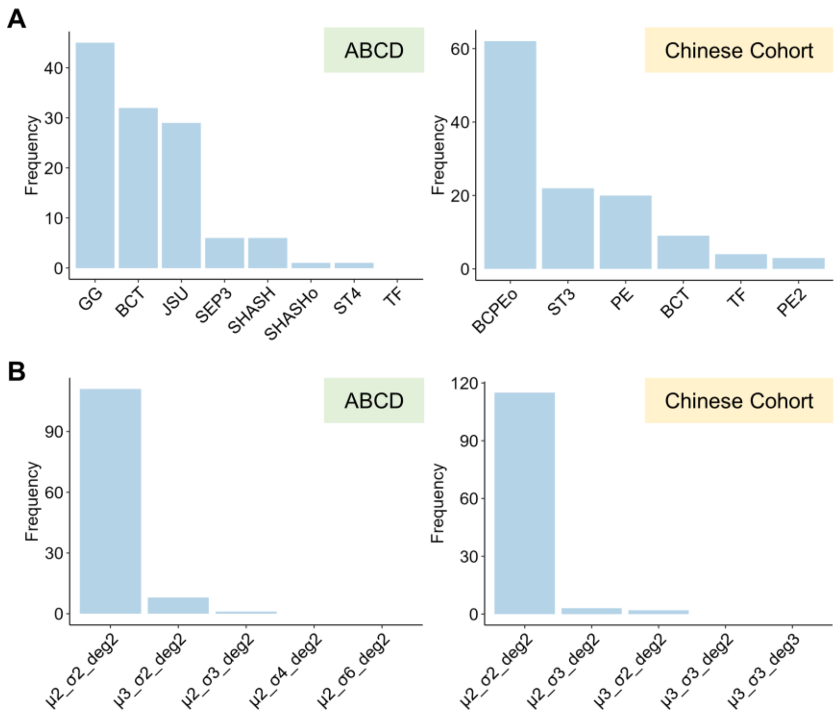


**Fig. S3 | Selection of optimal distribution and B-spline parameters for the normative models. A**, The selection frequency of the optimal distribution for the GAMLSS models. For each of the 120 connections, the optimal distribution was selected based on the lowest Bayesian Information Criterion (BIC). This panel displays the selection frequency for distributions that successfully converged across all models. The generalized gamma (GG) distribution was most frequently selected for the ABCD dataset, while the Box–Cox Power Exponential (BCPEo) distribution was selected for the Chinese Cohort. Only the frequency of distributions which converged across all 120 connectional models was presented. **B**, The selection frequency of the optimal parameters for the B-spline function used to model age. Similar to the distribution selection, the optimal parameter set (degrees of freedom and polynomial degree) was determined for each of the 120 connections based on the lowest BIC. A degree of freedom (df) of 2 for location ($\mu$) and scale ($\sigma$) and a polynomial degree of 2 was the most frequently selected parameter set for both datasets. The panel presents the top six most frequently selected B-spline parameter sets.

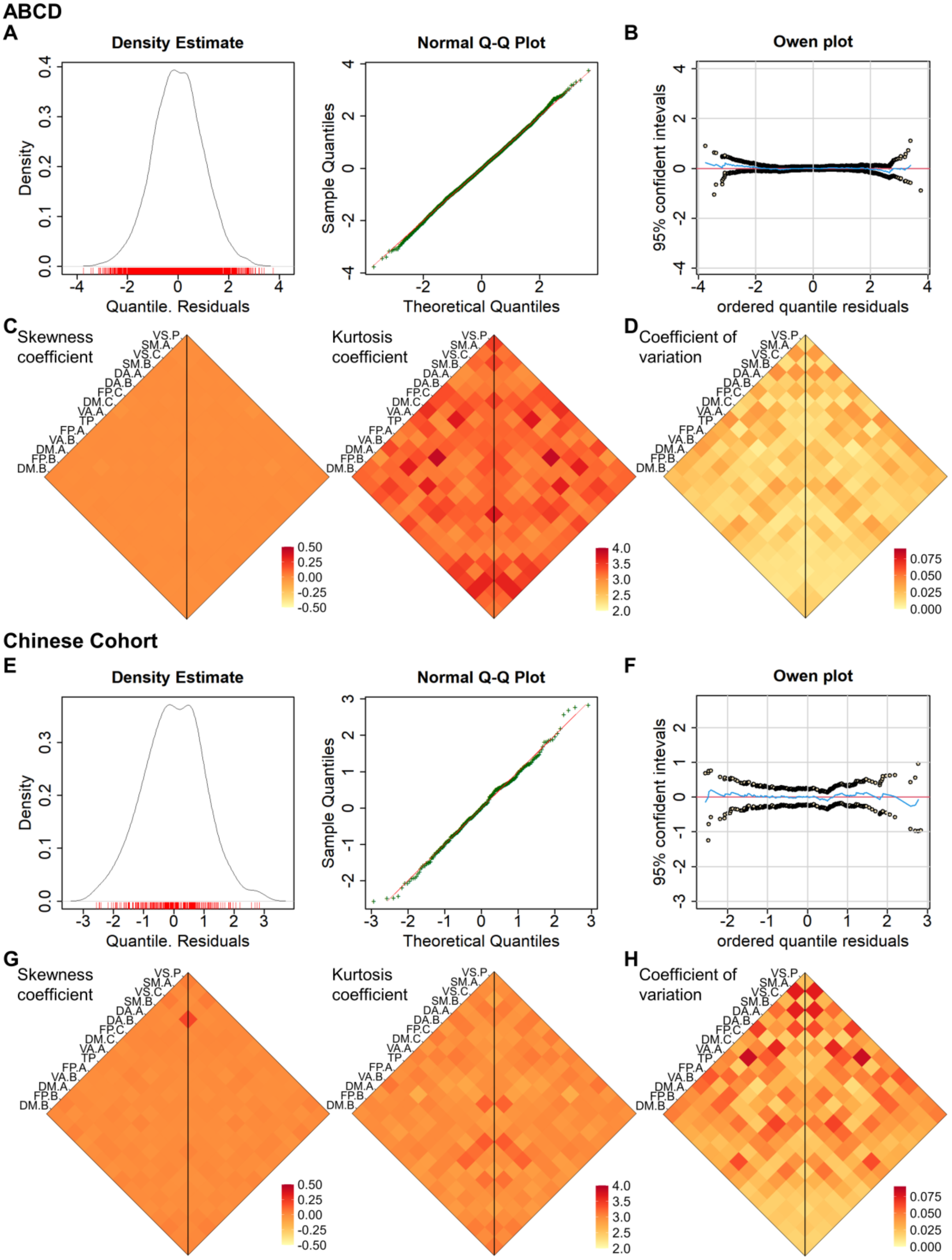


**Fig. S4 | Goodness-of-fit and reliability of the normative models for each connection.** **A**,**E**, Distribution plots and Q-Q plots for the normalized quantile residuals from the (**A**) ABCD and (**E**) Chinese Cohorts. The close alignment of sample quantiles to the theoretical quantiles indicates that the residuals are approximately normally distributed. **B**,**F**,

Detrended transformed Owen's plots for the (**B**) ABCD and (**F**) Chinese Cohorts. The 95% confidence intervals around the residuals consistently include zero, indicating an acceptable model fit. **C**,**G**, Skewness and kurtosis coefficients of the normative quantile residuals for all 120 connection models from the (**C**) ABCD and (**G**) Chinese Cohorts. For both datasets, all models had skewness coefficients within -0.5 to 0.5 and excess kurtosis coefficients within 2 to 4, further supporting the approximate normality of the residuals. **D**,**H**, Results of a 1,000-iteration bootstrap analysis assessing model reliability for the (**D**) ABCD and (**H**) Chinese Cohorts. The plots show the mean coefficient of variation (CV) of the median developmental trajectories derived from the 1,000 bootstrapped models. For all connections, the average CV did not exceed 0.1, indicating good model reliability.

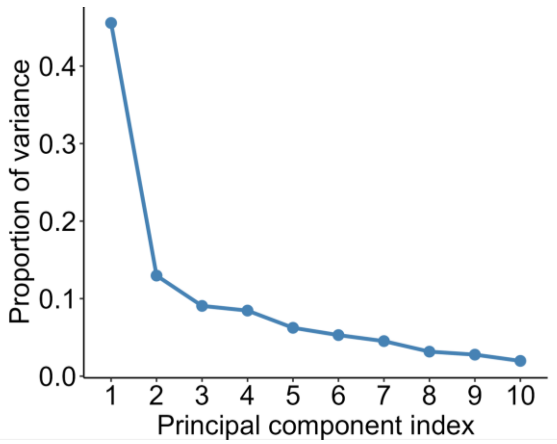


**Fig. S5 | Variance explained by principal components in children with ADHD from the Chinese cohort.** Proportion of variance explained by each principal component derived from SC deviations of the 10 connections that showed age-related effects replicated in the Chinese cohort.

**Table S1. Demographic and clinical symptoms of participants from the ABCD dataset.**

| | TD training | TD test | ADHD | $P_a$ | $P_b$ |
|---|---|---|---|---|---|
| N | 4599 | 2088 | 1114 | | |
| Visit (%) | | | | 0.577 | 0.001 |
| Baseline | 2177 (47.3) | 978 (46.8) | 597 (53.6) | | |
| 2-Year follow-up | 1984 (43.1) | 894 (42.8) | 401 (36.0) | | |
| 4-Year follow-up | 438 (9.5) | 216 (10.3) | 116 (10.4) | | |
| Sex = M (%) | 2285 (49.7) | 1038 (49.7) | 800 (71.8) | 1 | <0.001 |
| Age (mean (SD)) | 11.19 (1.47) | 11.23 (1.53) | 11.05 (1.52) | 0.413 | 0.003 |
| Mean FD (mean (SD)) | 0.61 (0.23) | 0.61 (0.23) | 0.65 (0.25) | 0.62 | <0.001 |
| Race/ethnicity (%) | | | | 0.139 | <0.001 |
| Asian | 106 (2.3) | 51 (2.4) | 8 (0.7) | | |
| Black | 586 (12.7) | 264 (12.6) | 117 (10.5) | | |
| Hispanic | 978 (21.3) | 397 (19.0) | 180 (16.2) | | |
| Other | 439 (9.5) | 183 (8.8) | 129 (11.6) | | |
| White | 2490 (54.1) | 1193 (57.1) | 680 (61.0) | | |
| Externalizing (mean (SD)) | 40.38 (7.44) | 40.58 (7.23) | 55.51 (10.18) | 0.283 | <0.001 |
| Attention problem (mean (SD)) | 51.31 (2.72) | 51.35 (2.81) | 63.46 (8.09) | 0.547 | <0.001 |
| ADHD problem (mean (SD)) | 51.03 (2.62) | 51.01 (2.57) | 62.72 (7.51) | 0.828 | <0.001 |
| Sites (%) | | | | - | - |
| Site01 | 148 (3.2) | 66 (3.2) | 22 (2.0) | | |
| Site02 | 295 (6.4) | 125 (6.0) | 58 (5.2) | | |
| Site03 | 273 (5.9) | 120 (5.7) | 45 (4.0) | | |
| Site04 | 319 (6.9) | 142 (6.8) | 90 (8.1) | | |
| Site05 | 158 (3.4) | 75 (3.6) | 32 (2.9) | | |
| Site06 | 198 (4.3) | 91 (4.4) | 88 (7.9) | | |
| Site07 | 73 (1.6) | 38 (1.8) | 32 (2.9) | | |
| Site08 | 97 (2.1) | 49 (2.3) | 38 (3.4) | | |
| Site09 | 162 (3.5) | 81 (3.9) | 26 (2.3) | | |
| Site10 | 296 (6.4) | 132 (6.3) | 60 (5.4) | | |
| Site11 | 144 (3.1) | 68 (3.3) | 55 (4.9) | | |
| Site12 | 233 (5.1) | 107 (5.1) | 78 (7.0) | | |
| Site13 | 263 (5.7) | 116 (5.6) | 71 (6.4) | | |
| Site14 | 302 (6.6) | 135 (6.5) | 51 (4.6) | | |
| Site15 | 136 (3.0) | 61 (2.9) | 24 (2.2) | | |
| Site16 | 425 (9.2) | 191 (9.1) | 105 (9.4) | | |
| Site17 | 225 (4.9) | 101 (4.8) | 46 (4.1) | | |
| Site18 | 120 (2.6) | 62 (3.0) | 39 (3.5) | | |
| Site19 | 182 (4.0) | 84 (4.0) | 36 (3.2) | | |
| Site20 | 303 (6.6) | 136 (6.5) | 66 (5.9) | | |
| Site21 | 247 (5.4) | 108 (5.2) | 52 (4.7) | | |

Note: The number of participants and percentages were displayed for categorical variables, while mean and standard deviation (SD) were displayed for numeric variables. Ages in months and site information were obtained from ‘abcd_y_lt.csv’. Ages were then converted to years by dividing by 12. Sex and race/ethnicity data were obtained from ‘abcd_p_demo.csv’. Symptom scores were derived from the parent-reported Child Behavior Checklist (CBCL), found in ‘mh_p_cbcl.csv’. Diagnoses were obtained from the parent-reported computerized version of the Kiddie Schedule for Affective Disorders and Schizophrenia (KSADS-COMP), available in ‘mh_p_ksads_ss.csv’. Statistical comparisons were performed using independent samples T-tests for numeric variables and chi-square tests for categorical variables. Specifically, $P_a$ denotes comparisons between the TD training and test sets, while $P_b$ represents comparisons between the TD test set and the ADHD group.

**Table S2. Demographic and clinical symptoms of participants from the Chinese Cohort.**

| Sample for validation analysis | | | |
|---|---|---|---|
| | TD | ADHD | *P* |
| N | 292 | 468 | |
| Sex = M (%) | 156 (53.4) | 367 (78.4) | <0.001 |
| Age (mean (SD)) | 10.10 (2.07) | 10.15 (2.05) | 0.773 |
| Mean FD (mean (SD)) | 0.55 (0.19) | 0.46 (0.23) | <0.001 |
| Study = PKU6 (%) | 97 (33.2) | 430 (91.9) | - |
| Sample for medication study (PKU6) | | | |
| | ADHD (ATX) | ADHD (MPH) | *P* |
| N | 40 | 72 | |
| Sex = M (%) | 35 (87.5) | 51 (70.8) | 0.077 |
| Age (mean (SD)) | 10.18 (2.02) | 10.22 (2.05) | 0.910 |
| Mean FD (mean (SD)) | 0.44 (0.26) | 0.40 (0.16) | 0.341 |
| Inattention-Pre (mean (SD)) | 18.48 (4.46) | 18.18 (4.59) | 0.743 |
| Hyperactivity-Pre (mean (SD)) | 12.25 (7.50) | 10.76 (6.63) | 0.280 |
| Inattention-Post (mean (SD)) | 11.18 (5.46) | 11.43 (5.47) | 0.813 |
| Hyperactivity-Post (mean (SD)) | 7.50 (5.79) | 5.94 (4.74) | 0.128 |

Note: The number of participants and percentages were displayed for categorical variables, while mean and SD were displayed for numeric variables. Statistical comparisons were performed using independent samples T-tests for numeric variables and chi-square tests for categorical variables. ATX: atomoxetine; MPH: methylphenidate; Pre: pre-treatment; Post: post-treatment.

**Table S3. Interaction effects between age and diagnosis on SC deviations in the ABCD dataset.**

| Label | $F_{ADHD}$ | $P_{FDR, ADHD}$ | $F_{TD}$ | $P_{FDR, TD}$ | $\chi^2_{Iteraction}$ | $P_{FDR, Interaction}$ |
|---|---|---|---|---|---|---|
| **SM.A-SM.B** | 7.68 | 0.008 | 0.56 | 0.752 | 8.64 | 0.042 |
| **SM.A-VA.B** | 6.89 | 0.014 | 0.39 | 0.837 | 8.43 | 0.042 |
| **SM.A-DM.A** | 4.89 | 0.043 | 3.17 | 0.304 | 8.23 | 0.043 |
| SM.A-FP.B | 8.29 | 0.007 | 6.05 | 0.063 | 4.66 | 0.126 |
| **SM.A-DM.B** | 8.96 | 0.006 | 3.59 | 0.273 | 7.83 | 0.044 |
| SM.B-DM.A | 5.48 | 0.029 | 0.98 | 0.625 | 3.04 | 0.230 |
| **DA.A-FP.C** | 5.84 | 0.023 | 1.19 | 0.584 | 11.29 | 0.037 |
| DA.A-TP | 4.67 | 0.043 | 0.61 | 0.729 | 3.80 | 0.177 |
| **DA.A-DM.B** | 13.87 | 0.003 | 4.19 | 0.234 | 9.00 | 0.042 |
| **DA.B-FP.A** | 5.83 | 0.023 | 0.30 | 0.885 | 7.27 | 0.046 |
| DA.B-FP.B | 7.83 | 0.008 | 5.10 | 0.156 | 3.68 | 0.181 |
| DA.B-DM.B | 5.32 | 0.041 | 3.77 | 0.273 | 6.58 | 0.060 |
| **FP.C-TP** | 10.08 | 0.003 | 0.25 | 0.889 | 8.85 | 0.042 |
| FP.C-FP.A | 5.64 | 0.023 | 2.77 | 0.379 | 6.27 | 0.069 |
| FP.C-DM.A | 5.99 | 0.023 | 1.00 | 0.625 | 5.54 | 0.100 |
| **FP.C-DM.B** | 12.77 | 0.003 | 1.22 | 0.584 | 11.13 | 0.037 |
| DM.C-TP | 7.24 | 0.008 | 1.47 | 0.545 | 5.02 | 0.111 |
| DM.C-FP.B | 4.82 | 0.041 | 3.38 | 0.280 | 1.97 | 0.368 |
| **FP.A-VA.B** | 8.43 | 0.003 | 2.15 | 0.481 | 8.17 | 0.043 |
| **FP.A-FP.B** | 9.10 | 0.003 | 1.44 | 0.545 | 8.86 | 0.042 |
| **FP.A-DM.B** | 11.73 | 0.003 | 2.72 | 0.383 | 8.19 | 0.042 |
| **VA.B-DM.B** | 9.92 | 0.003 | 2.57 | 0.383 | 14.26 | 0.017 |
| DM.A-FP.B | 6.12 | 0.023 | 1.68 | 0.543 | 3.39 | 0.205 |
| **FP.B-DM.B** | 7.22 | 0.007 | 1.63 | 0.543 | 7.40 | 0.046 |
| DM.B-DM.B | 5.02 | 0.039 | 2.42 | 0.385 | 5.01 | 0.112 |

Note: *F* statistics were reported for the nonlinear age effects on SC deviations in the TD and ADHD groups separately. FDR correction was applied across all 120 edges for each group. For the interaction effects between age and diagnosis on SC deviations, we reported the $\chi^2$ statistics. The *P* values of the interaction effects were corrected across the 25 edges that showed significant age effects in the ADHD group. All 25 edges with significant age effects in the ADHD group are presented in the table, while only the labels of the 14 edges showing ADHD-specific effects ($P_{FDR, interaction} < 0.05$) are highlighted in bold.

**Table S4. Age effects on SC deviations replicated in the children with ADHD in Chinese cohort.**

| Label | $F_{ADHD}$ | $P_{FDR, ADHD}$ |
|---|---|---|
| **SM.A-SM.B** | 8.87 | 0.001 |
| SM.A-VA.B | 0.36 | 0.698 |
| SM.A-DM.A | 1.62 | 0.262 |
| **SM.A-DM.B** | 4.78 | 0.017 |
| DA.A-FP.C | 0.66 | 0.552 |
| **DA.A-DM.B** | 14.01 | 0.001 |
| **DA.B-FP.A** | 4.91 | 0.013 |
| **FP.C-TP** | 9.08 | 0.001 |
| FP.C-DM.B | 3.24 | 0.056 |
| **FP.A-VA.B** | 9.88 | 0.001 |
| **FP.A-FP.B** | 6.65 | 0.002 |
| **FP.A-DM.B** | 10.46 | 0.001 |
| VA.B-DM.B | 1.31 | 0.319 |
| **FP.B-DM.B** | 11.54 | 0.001 |

Note: *F* statistics were reported for the nonlinear age effects on SC deviations in the ADHD group. FDR correction was applied across all the 14 edges, and the labels of edges with significant age effects ($P_{FDR} < 0.05$) are shown in bold.

**Table S5. Image acquisition parameters for T1-weighted images and diffusion MRI for each dataset.**

| | Sequence | TR (ms) | TE (ms) | TI (ms) | Flip angle (°) | FOV ($mm^2$) | Slices | Voxel Size (mm) | Diffusion directions | b-values ($s/mm^2$) |
|---|---|---|---|---|---|---|---|---|---|---|
| ABCD (3T SIEMENS) | | | | | | | | | | |
| T1WI | MPRAGE | 2500 | 2.88 | 1060 | 8 | 256×256 | 176 | 1.0 | NA | NA |
| dMRI (PA) | Multiband EPI | 4100 | 88 | NA | 90 | 240×240 | 81 | 1.7 | 96 | 500, 1000, 2000, 3000 |
| fmap (AP) | Single-band EPI | 12400 | 89 | NA | 90 | 240×240 | 81 | 1.7 | 0 | 0 |
| ABCD (3T Philips) | | | | | | | | | | |
| T1WI | 3D TFE | 6.31 | 2.9 | 1060 | 8 | 256×240 | 225 | 1.0 | NA | NA |
| dMRI (PA) | Multiband EPI | 5300 | 89 | NA | 78 | 240×240 | 81 | 1.7 | 96 | 500, 1000, 2000, 3000 |
| fmap (AP) | Multiband EPI | 5300 | 89 | NA | 78 | 240×240 | 81 | 1.7 | 0 | 0 |
| ABCD (3T GE) | | | | | | | | | | |
| T1WI | MPRAGE | 2500 | 2 | 1060 | 8 | 256×256 | 208 | 1.0 | NA | NA |
| dMRI (PA) | Multiband EPI | 4100 | 81.9 | NA | 77 | 240×240 | 81 | 1.7 | 96 | 500, 1000, 2000, 3000 |
| fmap (AP) | Multiband EPI | 4100 | 81.9 | NA | 77 | 240×240 | 81 | 1.7 | 0 | 0 |
| Chinese Cohort-EFNY (3T SIEMENS Prisma with 64-channel head coil) | | | | | | | | | | |
| T1WI | MPRAGE | 1500 | 1.87 | 756 | 10 | 256×256 | 208 | 0.8 | NA | NA |
| dMRI (PA) | Multiband EPI | 3100 | 86 | NA | 90 | 205×205 | 84 | 1.8 | 120 | 500, 1000, 2000, 3000 |
| fmap (AP) | Multiband EPI | 3100 | 86 | NA | 90 | 205×205 | 84 | 1.8 | 0 | 0 |
| Chinese Cohort-PKU6 (3T GE discovery MR750 with 8-channel head coil) | | | | | | | | | | |
| T1WI | 3D SPGR | 6700 | 2.9 | 450 | 12 | 256×256 | 192 | 1.0 | NA | NA |
| dMRI (PA) | Single-band EPI | 8980 | 92 | NA | 90 | 240×240 | 70 | 2.0 | 64 | 1000 |

Note: T1WI: T1-weighted imaging; dMRI: diffusion magnetic resonance imaging; MPRAGE: magnetization prepared rapid gradient echo; TFE: Turbo/fast Field Echo; EPI: echo planar imaging; 3D SPGR: three-dimensional spoiled gradient recalled; AP: anterior-to-posterior phase-encoding direction; PA: posterior-to-anterior phase-encoding direction.

**Table S6. GAMLSS candidate continuous distributions with three or four parameters.**

| Distribution | R Name | $\mu$ | $\sigma$ | $\nu$ | $\tau$ |
|---|---|---|---|---|---|
| Box-Cox Cole and Green | BCCG() | identity | log | identity | - |
| Box-Cox Power Exponential | BCPE() | identity | log | identity | log |
| Box-Cox Power Exponential (original/link variant) | BCPEo() | log | log | identity | log |
| Box-Cox-t | BCT() | identity | log | identity | log |
| Exponential Gaussian | exGAUS() | identity | log | log | - |
| Exponential generalized Beta type 2 | EGB2() | identity | log | log | log |
| Generalized Beta type 2 | GB2() | log | log | log | log |
| Generalized Gamma | GG() | log | log | identity | - |
| Generalized Inverse Gaussian | GIG() | log | log | identity | - |
| Generalized t | GT() | identity | log | log | log |
| Johnson's SU | JSU() | identity | log | identity | log |
| Power Exponential | PE() | identity | log | log | - |
| Power Exponential type 2 | PE2() | identity | log | log | - |
| Skew Power Exponential type 1 | SEP1() | identity | log | identity | log |
| Skew Power Exponential type 2 | SEP2() | identity | log | identity | log |
| Skew Power Exponential type 3 | SEP3() | identity | log | log | log |
| Skew Power Exponential type 4 | SEP4() | identity | log | log | log |
| Sinh-Arcsinh | SHASH() | identity | log | log | log |
| Sinh-Arcsinh (original Jones-Pewsey form) | SHASHo() | identity | log | identity | log |
| Skew t type 1 | ST1() | identity | log | identity | log |
| Skew t type 2 | ST2() | identity | log | identity | log |
| Skew t type 3 | ST3() | identity | log | log | log |
| Skew t type 4 | ST4() | identity | log | log | log |
| Skew t type 5 | ST5() | identity | log | identity | log |
| t-distribution | TF() | identity | log | log | - |

Note: The default link functions were annotated.
